# Primary productivity in subsidized green-brown food webs

**DOI:** 10.1101/2021.12.22.473860

**Authors:** Yuval R. Zelnik, Stefano Manzoni, Riccardo Bommarco

## Abstract

Ecosystems worldwide receive large amounts of nutrients from both natural processes and human activities. While direct subsidy effects on primary productivity are relatively well known (the green food web), the indirect effects of subsidies on producers as mediated by the brown food web and predators have been neglected. With a dynamical green-brown food web model, parameterized using empirical estimates from the literature, we illustrate the effect of nutrient subsidies on net primary productivity (i.e., after removing loss to herbivory) in two generic ecosystems, terrestrial and aquatic. We find that nutrient subsidies increase net primary productivity because more nutrients are available, but this effect saturates with higher subsidies. Changing the quality of subsidies from inorganic to organic tends to increase net primary productivity in terrestrial ecosystems, but less often so in aquatic ecosystems. This occurs when organic nutrient inputs promote detritivores in the brown food web, and hence predators that in turn control herbivores, thus promoting primary productivity. This previously largely overlooked effect is further enhanced by ecosystem properties such as fast decomposition and low rates of nutrient additions, and demonstrates the importance of nutrient subsidy quality on ecosystem functioning.

## 2 Introduction

No ecosystem is an island, entirely cut off from external influences. On the contrary, the influx of materials and organisms into ecosystems, known as allochthonous inputs or subsidies, have a significant impact on the state and functioning of ecosystems (Polis *et al*., 1997; Palumbi, 2003). The subsidies can take various forms, including material passively moved by water and air (Bobbink *et al*., 2010), animals that move across ecosystems for various activities such as foraging (Marczak *et al*., 2007; Buendía *et al*., 2018), and, an increasingly important phenomenon for ecosystems world wide, input of resources such as nutrients through human activities (Newsome *et al*., 2015; Raun & Johnson, 1999). Subsidies can have diverse impacts. Adding limiting nutrients often increases overall primary production (Polis *et al*., 1997; Montagano *et al*., 2018), but it can also decline if, for instance, the added nutrients promote plant-microbe competition for other nutrients (Čapek *et al*., 2018). Introducing consumers changes the fluxes in the ecosystem and thereby affects biomass distributions across trophic levels (Allen & Wesner, 2016), and introduced species can change the behavior of the ecosystem entirely, as it creates novel interactions and pathways within the ecosystem (Baxter *et al*., 2004).

Subsidies can move both horizontally between food web compartments at a certain trophic level and vertically across trophic levels within a food web. Horizontal transfer occurs due to nutrient flow and interaction of organisms between food webs, and can occur through physical transfer of material between spatially separated webs in an ecosystem, or through species interactions in linked food webs (Nakano & Murakami, 2001; Baxter *et al*., 2004). An example of the latter is consumer species in green-brown food webs that rely on both primary producers in the green channel and detrital matter in the brown channel (Moore *et al*., 2004; Allen & Wesner, 2016). Subsidies also lead to vertical exchanges between trophic levels within the food web. Subsidization from below, by influx of nutrients or direct subsidization of primary producers, propagates up to consumer species, increasing the overall biomass and the shape of biomass distribution in the food web (Hines *et al*., 2006). Input of predators leads to pressure on the herbivores they consume, and can lead to trophic cascades where primary producers are released from herbivory pressure due to predation (Leroux & Loreau, 2008; Newsome *et al*., 2015; Galiana *et al*., 2021). Hence, subsidies can cause positive albeit indirect effects on primary production via trophic interactions, but outcomes of such combined horizontal and vertical effects of subsidies in green-brown food webs are poorly explored.

Research on the impact of subsidies has centered primarily on either the horizontal or the vertical flows, without bringing these two perspectives together (Rooney *et al*., 2006). For instance, landscape ecologists have often investigated the horizontal transfer between habitats (Darimont *et al*., 2009), while food web ecologists have focused on vertical flows that can change biomass distributions and can lead to trophic cascades (Leroux & Loreau, 2008). However, flow of energy and nutrients due to subsidies can be more nuanced due to the complexity of interactions within ecosystems, intertwining the horizontal and vertical flows. Research on subsidies has largely focused on green food webs (Nakano & Murakami, 2001; Loreau & Holt, 2004; Bobbink *et al*., 2010), with more recent interest in brown food webs (Allen & Wesner, 2016; McCary *et al*., 2020), but linking the two in integrated green-brown food webs has only just begun.

Nutrient subsidies into the brown channel of the food web has recently been show to impact plant productivity (Riggi & Bommarco, 2019; Aguilera *et al*., 2021): the subsidies provide food for detritivores, which in turn increases predator populations, and leads to a trophic cascade where plants grow more as herbivory pressure is reduced by predators. However, a theoretical understanding of these processes and their relevance has lagged behind. While some theoretical investigations have been done on how organic matter subsidies affect primary producers and nutrient recycling (Leroux & Loreau, 2008; Gounand *et al*., 2014; Spiecker *et al*., 2016), we know of only two studies in which the full green-brown food web was considered in the context of subsidies (Attayde & Ripa, 2008; McCary *et al*., 2020). However, they did not consider how the addition rate and quality of nutrient subsidies affect the food web.

Here we develop and use a dynamical model to examine how nutrient subsidies affect primary production in coupled green-brown food webs in terrestrial and aquatic ecosystems. We focus on nitrogen as a key nutrient whose cycle has been widely disrupted by human activities, and measure net primary productivity, the productivity remaining after herbivory, as a function of nutrient subsidy. The nutrient subsidy properties we consider are the amounts of inorganic and organic nitrogen in detrital matter (e.g., green or animal manure), which are consumed by the green and brown channels of the ecosystem, respectively, and relative mixtures of these. Combining the model with data from the literature on nitrogen flows and stocks we aim to assess: 1) how the strength and quality of subsidies affect net primary productivity, i.e., accounting for herbivory, and 2) how the defining food web properties of consumption, nutrient conversion and metabolic rates for consumers, and ecosystem properties such as mineralization rate and primary producer mortality, modify the subsidy effect on primary producers in two ecosystem types, aquatic and terrestrial.

## 3 Methods

### 3.1 Dynamical Model

To model the dynamics of subsidized green-brown food webs, we write six mass balance equations describing the changes in nitrogen stock of four different functional groups of organisms (primary producers and faunal groups) and two substrate compartments (organic and inorganic nitrogen) (Fig. 1). While we focus on nitrogen as a key limiting nutrient in most ecosystems, we keep the model general, to open for the possibility to describe the dynamics of other nutrients as well. In order to keep the model simple and generic, we follow Barbier & Loreau (2019) in defining the food web interactions, using a type *I* functional response for consumption terms, together with self-regulation (Barabás *et al*., 2017) of each species compartment. A conceptual diagram is given in Fig. 1, and the model’s dynamics are detailed by the coupled ordinary differential equations, given in eqs. 1.

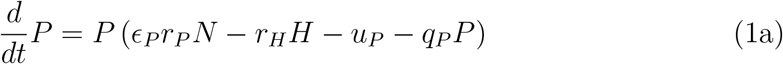

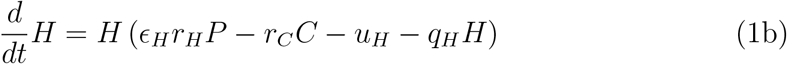

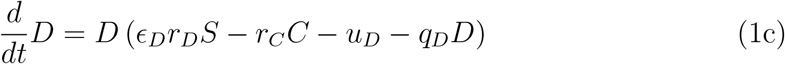

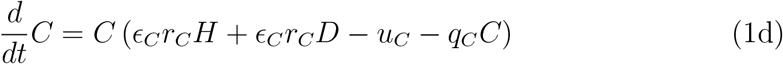

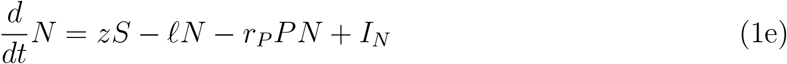

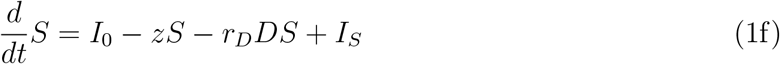

**Figure 1:**
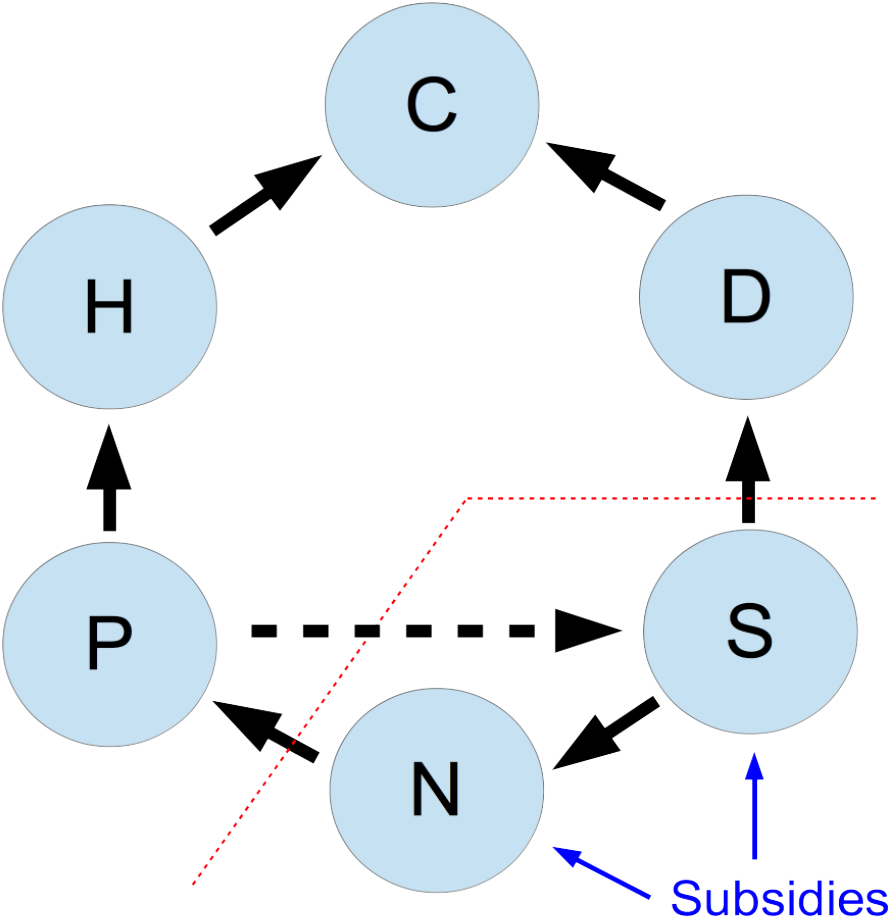
Interactions between the six compartments of an ecosystem model with nutrient subsidies. Compartments are: primary producers (**P**), herbivores (**H**), detritivores (**D**), predators (**C**), inorganic nutrients (**N**), active organic nitrogen (**S**). Black lines show nutrient flows between compartments; blue lines show the nutrient subsidies that are added to the ecosystem, the red line marks the separation between organism and substrate compartments, and the dashed arrow indicates nutrient recycling from dead primary producers to the organic nitrogen compartment.

Here, the six compartments are: P – primary producers, H – herbivores, D – detritivores (excluding microorganisms), C – predators, N – inorganic nitrogen, S – active organic nitrogen, which includes microbial decomposers. For each of the first four compartments (with *i* designating the specific compartment), the parameter *r_i_* represents the consumption coefficient of the trophic level below (in units of [*m*^2^(*gN*)^−1^*yr*^−1^]), *ϵ_i_* is the non-dimensional nutrient conversion efficiency associated with this consumption, *u_i_* is the natural mortality rate (units of [*yr*^−1^]), and *q_i_* is the self-regulation coefficient (e.g., light limitation for primary producers, or intra-guild predation for predators, in units of [*m*^2^(*gN*)^−1^*yr*^−1^]). In the two abiotic compartments, *z* is the rate of nitrogen mineralization, *ℓ* is the leaching/loss rate of inorganic nitrogen, both in units of [*yr*^−1^]. *I_N_* and *I_S_* are the influx rates of nitrogen into the *N* and *S* compartments, respectively, which are interpreted as subsidy fluxes of inorganic and organic nitrogen. We define the nutrient subsidy strength as *I* = *I_N_* + *I_S_*. The fraction of organic subsidy, which we call subsidy quality or *ψ*, is calculated as *I_S_/I*. Thus, *ψ* = 0 for strictly inorganic subsidy and *ψ* = 1 for only organic inputs. *I*_0_ is the input of nutrients from recycled materials from primary producers, given as *I*_0_ = *yu_P_P*, with *y* the recycling rate (the quantitatively smaller recycling from other compartments is neglected for simplicity).

We note in particular two modeling choices. By having self-regulation terms we can expect more well behaved dynamics (Barabás *et al*., 2017), e.g., avoiding oscillatory dynamics observed in previous studies (Attayde & Ripa, 2008). We focus on a type-I functional response, as it is simpler to analyze, and it is not a priori clear which type of functional response is the most relevant at the ecosystem level. Beyond type-I functional response, type-II functional response is often used in consumer-resource interactions (Attayde & Ripa, 2008; Wollrab *et al*., 2012), although its relevance has also been contested Jonsson (2017). Finally, we note that previous work (Wollrab *et al*., 2012) has often found that the specific type of functional response does not alter the qualitative repercussions of nutrient enrichment. We test in the SI a model with a type-II functional response, and conclude that a type-I functional response is more relevant for our modeling approach, as it is less likely to lead to extinct functional groups (e.g., no herbivores in the ecosystem) and temporal oscillations.

Nutrient recycling (*I*_0_), represented by the dashed horizontal arrow in Fig. 1, is a significant nutrient source in many natural ecosystems, especially terrestrial and macrophyte-dominated aquatic ecosystems (Manzoni et al., 2018). As we show in the Supporting Information (SI), including this process largely amounts to an increase in subsidy strength *I* and a rescaling of the subsidy’s organic fraction *ψ*. This effect of internal recycling can be explained by noticing that even with only the inorganic subsidy, the organic nutrient compartment is fed by the nutrient recycling of primary producer and animal necromass. Therefore, without loss of generality, we focus in the main text on ecosystem dynamics without nutrient recycling, by setting *y* = 0. This assumption allows for a simpler analysis and presentation, while retaining the qualitative behavior of the model (see the SI for qualitatively similar results where we include detritus recycling).

### 3.2 Model parameterization

To analyse the effects of subsidies on primary production, we parameterize the model for two generic ecosystem types, terrestrial and aquatic. These idealized ecosystems are used as baselines for analyses where ecosystem properties are varied widely, to ensure that results can be generalized. We base the model parameterization largely on a combined dataset from two previous data collections, which contain estimates of the nitrogen stocks and flow rates described in the model (Cebrian, 1999, 2004). We also use rough estimates of ecosystem properties such as nutrient conversion efficiencies from a dozen additional studies, detailed in the Data collection subsection below. All values are converted to units of grams of nitrogen per square meter for stocks, and per year for rates. As explained below in the Parameter derivation subsection, we assume that eqs. 1 are in equilibrium, and use the empirically-derived stock and rate estimates to back-calculate the model parameters for the terrestrial and aquatic ecosystem, respectively. We do this by estimating the production fluxes of the four compartments (*P,H,D,C*), and using these flux values together with values for fraction of production to estimate the various parameters. Finally, as explained in the Parameter exploration subsection, we use two methods to numerically explore the consequences on primary production of a wide range of possible parameter values representing different ecosystems.

### 3.3 Data collection

We combine the datasets from Cebrian (1999, 2004) according to ecosystem type: terrestrial and aquatic. Terrestrial ecosystems include those labelled as forests, grasslands, savannas, drylands and boreal ecosystems. Aquatic ecosystems include all marine and freshwater ecosystems, such as oceans, seagrass meadows, coral reefs, and lakes. We exclude data for marshes and swamps from either ecosystem type. With these definitions, we obtain 210 records for the terrestrial and 534 records for the aquatic ecosystem. The data types for each record (and their associated units) are: primary producer nitrogen content (*ν_P_* [*gN/gDW*]; *DW* stands for dry weight), primary producer stocks (*B_P_*[*gCm*^−2^]), herbivore stocks (*B_H_*[*gCm*^−2^]), mineralization rate (*z*[*yr*^−1^]), decomposition flux (*f_D_*[*gCyr*^−1^*m*^−2^]), detrital production (*f_M_*[*gCyr*^−1^*m*^−2^]), primary production (*f_P_*[*gCyr*^−1^*m*^−2^]), herbivory fraction (*ρ_PH_*). The geometric average of the available values is calculated for each data type and used in the subsequent estimation of the model parameters for each ecosystem type. The use of a geometric average is appropriate due to the wide range of values for various ecosystems Bar-On *et al*. (2018). We use the primary producer average nitrogen content, together with a ratio of carbon to dry-weight ratio of 2.5 (Cebrian & Lartigue, 2004), to convert *B_P_* and all fluxes from carbon to nitrogen, for instance *P* = *B_P_* · (*ν_P_*/2.5). We similarly use a carbon to nitrogen ratio for animals of 5 (Allgeier *et al*., 2020), to convert *B_H_* to nitrogen stocks.

Beyond the data from Cebrian (1999, 2004), other estimations of ratios of stocks and process rates are necessary in order to parameterize our model. In absence of a cohesive dataset containing the required information for both terrestrial and aquatic ecosystems, we use representative information from a variety of sources. In doing do, we strive to define plausible values for the model parameters, so that nitrogen stocks and fluxes remain within reasonable ranges observed in nature.

We start by estimating the ratios between compartment stocks, i.e., of predators to herbivores *σ_CH_*, detritivores to herbivores *σ_DH_*, and of inorganic to organic nitrogen *σ_NS_*. From a global estimation of biomass in the oceans (Bar-On *et al*., 2018), we see that the biomass of fish and large invertebrates is roughly similar to, or slightly smaller than, that of arthropods and protists, suggesting a similar biomass of herbivores (mainly plankton, e.g., protists), predators (mainly vertebrates, e.g., fish) and detritivores (all categories). We therefore assume that their stocks in the aquatic ecosystem to be the same, *σ_CH_* = *σ_DH_* = 1. Two studies reported biomass in different trophic groups in forests (Brockie & Moeed, 1986) and grasslands (Perkins *et al*., 2018), and found that detritivore biomass is an order of magnitude higher than (forest) and similar (grassland) to that of herbivores. Therefore, for terrestrial ecosystems we choose an intermediate value of detritivore biomass as five times the biomass of herbivores; i.e., *σ_DH_* = 5. The same studies also find that predator biomass is substantially smaller than herbivore biomass. Hence, we assume for the terrestrial ecosystem that *σ_CH_* = 0.2. These ratios for aquatic and terrestrial ecosystems are also roughly consistent with results for predator-prey systems (Hatton *et al*., 2015). A study of nitrogen cycling in aquatic ecosystems (Berman & Bronk, 2003) reported ratios between inorganic and organic nitrogen that were both higher and lower than 1. Conversely, Groffman & Rosi-Marshall (2013) found consistently lower inorganic nitrogen levels compared with organic nitrogen in terrestrial ecosystems, even when only considering the active proportion of soil organic material (around 5% of total organic matter; Paul (2016)). Based on this evidence, we assume *σ_NS_* =1 for the aquatic and *σ_NS_* = 0.5 for the terrestrial ecosystems.

We also estimate fractions of production lost to predation and to self-regulation, which includes intra-guild predation. The fraction of secondary production lost to predation could be high, reaching 90% in forests (Hairston & Hairston, 1993). Because we deem this value to be extremely high, we choose a more moderate value, and assume that *ρ_HC_* = *ρ_HD_* = 0.5 for both the terrestrial and aquatic ecosystems. Estimations for how much production is lost to self-regulation are virtually nonexistent. It has been suggested that for predators, self-regulation and predation coefficients are roughly proportional (Polis *et al*., 1989; Galiana *et al*., 2021). Assuming self-regulation and predation coefficients as equal (*r_C_* = *q_C_*), renders self-regulation fractions of *ρ_CC_* = 1.0 for the aquatic and *ρ_CC_* = 0.067 for the terrestrial ecosystem (see calculation in the Parameter derivation subsection). However, these values appear extreme and we therefore choose more moderate values, setting *ρ_CC_* = 0.5 for the aquatic and *ρ_CC_* = 0.1 for the terrestrial ecosystem. In doing so, we keep a large difference in self-regulation between ecosystem types, as suggested by the predation rates, but refrain from letting these difference be too large, potentially overshadowing other differences between ecosystem types. For other compartments there are no direct means to estimate self-regulation for a representative ecosystem. However, self-regulation for primary producers, e.g., via light limitation, is likely more substantial than for herbivores and detritivores which are more limited by predators and detrital matter, respectively (Hairston *et al*., 1960). We therefore choose, for both the terrestrial and aquatic ecosystems, small values for self-regulation fractions among herbivores and detritivores *ρ_HH_* = *ρ_DD_* = 0.05, but higher values for primary producers, *ρ_PP_* = 0.2.

Finally, we estimate loss/leaching rates and nutrient conversion efficiencies. For the terrestrial ecosystem, the ratios between annual loss flux and stocks of inorganic nitrogen are between 0 and 2 (Groffman & Rosi-Marshall, 2013), so that we can assume *ℓ* = 1[*yr*^−1^]. For the aquatic ecosystem, data from multiple lakes (Hohener & Gachter, 1993) gives a median value of *ℓ* = 2[*yr*^−1^], assuming the relevant water column depth is 10[*m*] (Middelboe & Markager, 1997). Next, we estimate nitrogen conversion efficiencies (which are generally higher than carbon conversion efficiencies, but by definition never higher than 1). Previous modeling studies have assumed nutrient conversion efficiencies for animals of 0.8 (Zou *et al*., 2016), as well as 0.5 and 0.25 (Attayde & Ripa, 2008), and both of these assume no loss for primary producer uptake of inorganic nitrogen. Following this we assume for primary producers *ϵ_P_* = 1, while for animals an intermediate value of 0.5, i.e., *ϵ_H_* = *ϵ_D_* = *ϵ_C_* = 0.5.

### 3.4 Parameter derivation

The values of all model parameters are estimated by inverting the model equations (eq. 1) for nitrogen fluxes, after setting nitrogen stocks, flows, and other ecosystem properties described in the previous subsection (Table 1). This approach is repeated for the terrestrial and aquatic ecosystems. Since we need values for all six compartments to determine the parameter values, our estimations of stocks are complemented by the ratios between nitrogen stocks in different compartments: *C* = *σ_CH_H, D* = *σ_DH_H*, and *N* = *σ_NS_S*.

**Table 1:** Parameters values used for simulations. Columns of aquatic and terrestrial correspond to the parameter estimations from the literature, as detailed in the Methods. In the column labelled ‘wide-coexistence’ a parameter set similar to the terrestrial one is reported, but with values chosen to have a wide region of coexistence. *z* is the rate of nitrogen mineralization, *ℓ* is the loss rate of inorganic nitrogen. The 16 other parameters correspond to four compartments (*P,H,D,C*), noted in the following by *i. r_i_*. consumption coefficient of the trophic level below; *ϵ_i_*: nutrient conversion efficiency associated with consumption; *u_i_*: natural mortality rate; *q_i_* self-regulation coefficient.

|  | Parameter values in: |  |  |  |
| --- | --- | --- | --- | --- |
|  | units | aquatic | terrestrial | wide-coexistence |
| $r_P$ | $[m^2(gN)^{-1}yr^{-1}]$ | 14.6 | 0.487 | 0.2 |
| $r_H$ | $[m^2(gN)^{-1}yr^{-1}]$ | 36.0 | 0.368 | 0.5 |
| $r_D$ | $[m^2(gN)^{-1}yr^{-1}]$ | 36.0 | 0.368 | 0.5 |
| $r_C$ | $[m^2(gN)^{-1}yr^{-1}]$ | 56.6 | 24.8 | 10 |
| $u_P$ | $[yr^{-1}]$ | 9.44 | 0.520 | 0.5 |
| $u_H$ | $[yr^{-1}]$ | 9.00 | 0.477 | 0.5 |
| $u_D$ | $[yr^{-1}]$ | 11.6 | 0.255 | 0.1 |
| $u_C$ | $[yr^{-1}]$ | 4.50 | 6.44 | 5 |
| $q_P$ | $[m^2(gN)^{-1}yr^{-1}]$ | 1.89 | 0.0260 | 0.02 |
| $q_H$ | $[m^2(gN)^{-1}yr^{-1}]$ | 5.66 | 0.496 | 0.5 |
| $q_D$ | $[m^2(gN)^{-1}yr^{-1}]$ | 7.30 | 0.0530 | 0.05 |
| $q_C$ | $[m^2(gN)^{-1}yr^{-1}]$ | 28.3 | 37.2 | 15 |
| $\epsilon_P$ | | 1 | 1 | 1 |
| $\epsilon_H$ | | 0.5 | 0.5 | 0.5 |
| $\epsilon_D$ | | 0.5 | 0.5 | 0.5 |
| $\epsilon_C$ | | 0.5 | 0.5 | 0.5 |
| $z$ | $[yr^{-1}]$ | 8.40 | 0.970 | 0.5 |
| $\ell$ | $[yr^{-1}]$ | 2 | 1 | 1 |

The growth term in eq. 1a, *f_P_* = *ϵ_P_r_P_N*, is essentially the primary production in nitrogen units, so that we can find the consumption rate of nutrients by primary producers as:

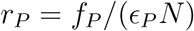

Similarly, the loss term due to herbivory is *r_H_H*, which can also be written as *f_PρPH_* (recall that *ρ_PH_* is the fraction of primary production consumed by herbivores). It follows that the consumption rate by herbivores is:

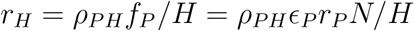

We also assume the detritivory rate to be the same as for herbivores, i.e., *r_D_* = *r_H_*, as both herbivores and detritivores are typically mobile animals feeding on sessile material. Finally, the loss term of herbivore biomass due to predation is *r_C_C*, which can also be written as *f_HρHC_*, where *ρ_HC_* is the fraction of herbivore production consumed by predators, and *f_H_* is the flux of nitrogen gained by herbivory. We thus find that:

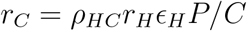

For primary producers, we use the flux detrital production *f_M_*, to find the natural mortality rate:

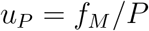

For the animal compartments we use the fraction left from predation to estimate the natural mortality:

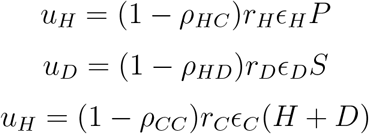

We similarly use the fraction of production lost to self-regulation, normalized by the compartment’s stock, to estimate the self-regulation coefficients:

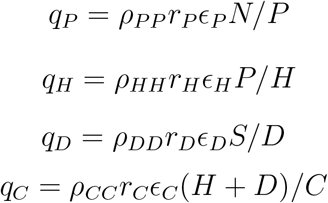

The values of the other parameters, namely the nutrient conversion efficiencies and the parameters describing losses from the organic nitrogen compartment (*z* and *ℓ*), are taken directly from their empirical estimates (Table 1). These derivations yield two parameter sets, of 18 parameters each, noted as the terrestrial and aquatic ecosystems in Table 1. In addition, we also construct an additional parameter set, based on the terrestrial parameters, noted as ‘wide coexistence’ in Table 1. The values in this parameter set are chosen to have a wide range of *ψ* values within the coexistence range (i.e., with all compartments extant), for *I* = 10[*gNm*^−2^*yr*^−1^], while still being very similar to the terrestrial ecosystem parameter set. While the choice of the ‘wide coexistence’ parameter values is arbitrary, it is done in order to visually demonstrate the effect of food web properties in Fig. 3. We test in the SI that the specific parameter value choice does not effect our qualitative results, and thus emphasize that our choices of this ‘wide coexistence’ parameter values are only for presentation purposes, and have no repercussions on our conclusions.

### 3.5 Simulations and parameter exploration

In all simulations and calculations we focus on the equilibrium of the ecosystem model. We find this equilibrium by integrating in time, until the maximal change in stocks per year, calculated for all nutrient input options, is less than 10^−^5, or until the simulation reached 1000 years (whichever comes first). We note that most simulations reach equilibrium in less than 100 years, and very few reach the 1000 years mark (Table S1.

To answer our questions on how 1) subsidy strength and quality affect net primary productivity (NPP) and 2) consumer rates and other ecosystem properties modify the subsidy effect on primary producers, we assess how NPP changes as the subsidy strength *I* and quality *ψ* are varied. The former is assumed to range between 1 and 100[*gNm*^−2^*yr*^−1^], and the latter across all possible values, between 0 (only inorganic nitrogen) and 1 (only organic nitrogen).

Besides varying the external inputs, we also explore the effect of other parameters: i) the effects of food web properties are assessed by changing several parameters in conjunction, and ii) an uncorrelated random parameter exploration of all parameters is performed to confirm the generality of the results.

First, we test the effect on ecosystem dynamics of three over-arching properties of the food web: consumption, conversion, and metabolic rates. We vary the baseline values of the parameters by a factor *γ*, so that the parameters are varied together in a perfectly coordinated manner. For consumption, we change consumption coefficients for the animal compartment *i* as: 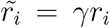. For conversion, we change nutrient conversion for the animal compartment *i* as 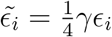 (the smaller range of nutrient conversion efficiencies is due to the constraint *ϵ_i_* ≤ 1). For metabolic rates, we change the consumption coefficients, natural mortality, and self-regulation coefficients in the three animal compartments as 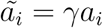, where *a* stands for *r, u* and *q*, and *i* identifies the animal compartment. In these analyses, we vary the control parameter *γ* on a logarithmic scale between 0.1 and 10. Our focus on variations in the properties of the animal compartments is motivated by our interest in how primary production is altered in the food web. However, we also test if concurrently changing the primary producer, *P*, parameters changes the qualitative results (see SI). To present the dependence of NPP on *ψ* (Fig. 3), we calculate the average value of NPP across the range of *I* values, and only show *ψ* values where compartments coexist, i.e., where all compartments have non-zero values, for at least some value of *I*. To present the NPP dependence on *I*, we do the same, except we average over the range of *ψ* values.

Second, to robustly test the effect of changing each parameter separately, we use a random parameter exploration. We take each of the 18 model parameters (all except *I* and *ψ*), and change their values relative to the baseline values of terrestrial or aquatic ecosystems (Table 1), without any correlation between parameters (in contrast to the previous analysis). For each parameter, we use a log-normal distribution, such that the base-10 logarithm values have a standard-deviation of 1, except for nutrient conversion efficiencies, for which we use a standard-deviation of 0.25 and we cap the values at 1. We repeat the sampling to create 20000 sets of randomly chosen parameters, and explore subsidy scenarios by considering a range of *ψ* values between 0 and 1, and seven values of *I* (1, 2, 5, 10, 20, 50, 100 [*gNm*^−2^*yr*^−1^]).

### 3.6 Ecosystem metrics for subsidy impact assessment

We focus on estimating net primary productivity (NPP), and its dependence on nutrient subsidies. The effect of herbivory is included in our definition of NPP, motivated by our aim to describe variations in ‘harvestable’ biomass increments, that is, primary producer biomass left after the herbivory fraction has been removed. This amounts to the term *u_p_P* in eq. 1, which is also directly proportional to the producer biomass *P* (as opposed to the term *r_P_ϵ_P_NP*, which includes the production lost to herbivory). Since productivity is strongly dependent on input levels *I*, and we change *I* across two orders of magnitude, it is useful to normalize NPP by *I*. We thus use a measure of normalized productivity, given by *u_P_P/I*. We note that this normalized productivity term is dimensionless, representing an ecosystem-level nitrogen use efficiency if we assume that mortality is due to harvesting in analogy to agricultural ecosystems (Scaini *et al*., 2020). As we show in the results, it is also useful examine how *P* changes when the subsidies are more organic, i.e., 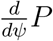 (*u_P_* and *I* are constant along the *ψ* axis, and we can therefore ignore them here). We normalize this term by the average primary producer biomass for a given level of subsidy strength *I*, and call this metric ‘Producer Sensitivity to Organic Subsidy’ 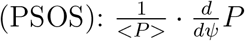.

## 4 Results

### 4.1 General responses of primary productivity to changes in subsidy strength and quality

We begin by looking at how subsidies, and in particular their strength *I* and their organic fraction *ψ*, affect net primary productivity (NPP) for our two representative ecosystems, terrestrial (left in Fig. 2) and aquatic (right). In each ecosystem, all parameters are constant except the subsidy parameters *I* and *ψ*. In general, NPP (given by the terms *u_P_P*) increases with larger subsidy strength, as more nutrients are converted into biomass. To highlight the effect of subsidies on the efficiency of nutrient conversion to biomass, in the following we focus on the normalized productivity *u_P_P/I*. When nitrogen inputs are low (the lower region in subsidy parameter space), normalized productivity is also low (dark blue). At high inputs, normalized productivity levels saturate as more biomass is lost to top consumers and to inefficiencies related to primary producer self-regulation (e.g., light limitation), as seen by the lower values of normalized productivity in Fig. 2.

**Figure 2:**
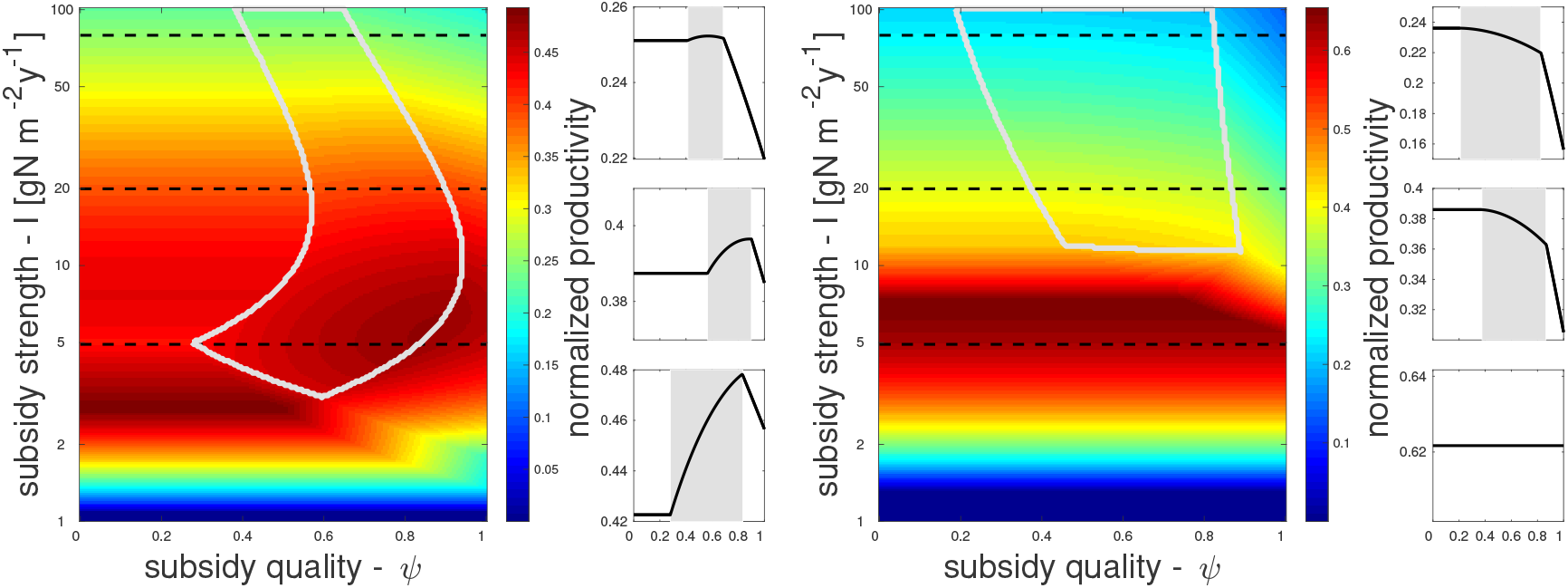
Normalized productivity *u_P_P/I* as a function of nitrogen subsidy strength (*I*) and quality (organic nitrogen fraction, *ψ*), in terrestrial (left) and aquatic (right) ecosystems. Besides *I* and *ψ*, other parameters are kept constant; gray lines enclose the coexistence region where all food web compartments (*P,H,D,C*) are extant. Small panels on the right of each parameter-space show normalized productivity at a constant value of *I* (gray background shows coexistence region), corresponding to the three horizontal dashed lines in each parameterspace.

As we increase the organic fraction of nitrogen inputs (moving left to right in the subsidy parameter space), NPP consistently decreases in the aquatic ecosystem, but within certain ranges of organic fraction it increases in the terrestrial one. This increase only occurs within the coexistence range where all compartments have nonzero values (marked by gray lines Fig. 2). The positive trend in the terrestrial ecosystem occurs due to strong predator control of herbivores allowing plants to grow more, as organic nitrogen subsidizes the detritivores, and thereby the predators. In contrast, decreases in NPP at higher organic fractions are due to larger amounts of nitrogen flowing into the brown food web instead of being used by primary producers.

The behavior outside the coexistence region can be explained as follows. When *D* = 0 (due to low *ψ*), net nitrogen mineralization equals the input of organic nitrogen (*zS* = *I_S_*), and hence at equilibrium all nitrogen inputs reach the *N* compartment, with *ψ* playing no further role. When *H* = 0 (due to high *ψ*) the only effect of detritivores on primary producers is competition over nitrogen, leading to a decrease of *P* with *ψ*, as the detritivores have opportunities to consume organic nitrogen before the primary producers can. Finally, when *C* = 0 (low I) no top-down control of the herbivores can occur, and hence *D* only exerts a negative effect via resource competition, similarly to the case where *H* = 0. Hereafter, we will focus on how nutrient subsidies determine NPP within the coexistence region, which changes depending on parameter selection, since we expect the species groups represented in our model – primary producers, herbivores, predators and detritivores – to all be extant in most ecosystems.

### 4.2 Effects of food web properties on productivity-subsidy relations

Focusing on the coexistence region, we can now assess how food web properties affect the link between subsidy strength and quality, and net primary productivity (NPP). We consider three food web properties encoded in parameters describing the three animal compartments (*H, D, C*): consumption, conversion, and metabolic rates.

NPP, normalized by subsidy strength, is affected jointly by subsidy quality *ψ* (top of Fig. 3) and strength *I* (bottom of Fig. 3), and by food web properties (indicated by different colors). This is shown for a specific set of initial ecosystem parameters (denoted as ‘wide-coexistence’ in Table 1), but we conduct a sensitivity analysis to test that the reported trends are consistent for a wide range of parameter sets, including parameters for aquatic ecosystems, as well as with internal recycling (see SI). Responses of normalized productivity are stronger, and often hump-shaped with a maximum at intermediate *I*, when subsidy strength is varied, compared with varying subsidy quality.

**Figure 3:**
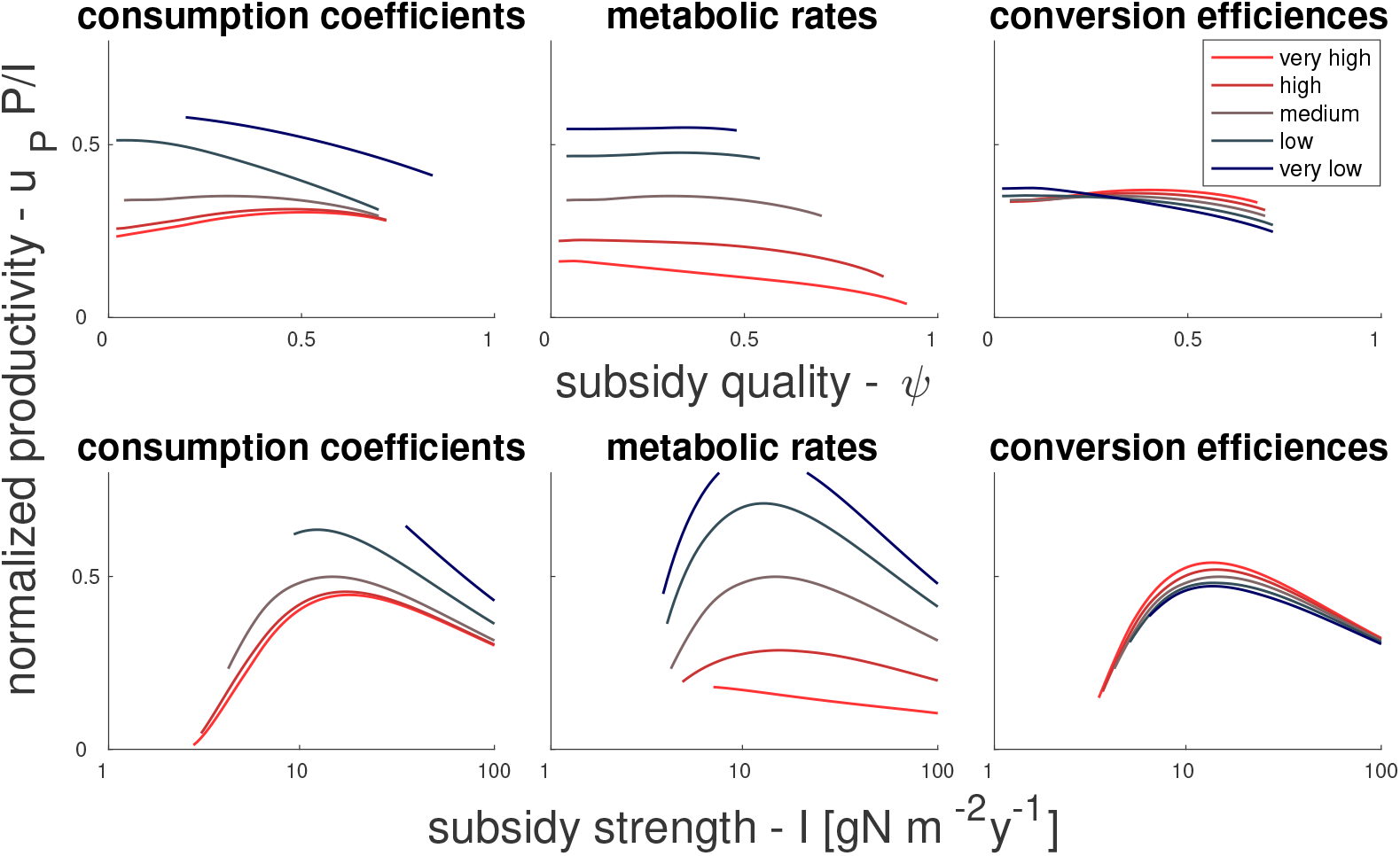
Effect of food web properties and subsidies on net primary productivity. Top (bottom) panels show normalized productivity as a function of subsidy quality (strength). Left, middle and right panels show sensitivities to food web properties as described by the parameters characterizing the animal compartments *H*, *D* and *C* (left: consumption coefficients; middle: consumption coefficients, mortality rates and self-regulation coefficients; right: nutrient conversion efficiencies:); lighter shades correspond to increasing values of model parameters. These changes are done relative to baseline parameters defined in Table 1 (column labelled ‘wide-coexistence’), corresponding to a terrestrial ecosystem with a large coexistence range. Results for other baseline parameter values are summarized in Fig. S10.

The interactions of food web properties with subsidies strength and quality cause a range of responses of net primary productivity. NPP decreases mildly with increasing consumption coefficients and strongly with increasing metabolic rates. In contrast, NPP varies less (and generally increases) with increasing nutrient conversion efficiency. Increasing metabolic rates causes a shift from mild positive dependency of NPP on subsidy quality to a mild negative dependency, whereas trends turn to positive when increasing consumption or conversion. The most notable interaction of food web properties with subsidy strength occurs when increasing metabolic rate, which causes a shift from hump-shaped to weakly negative relation between NPP and subsidy strength.

### 4.3 Analytical exploration of ecosystem processes driving productivity-subsidy relations

As we have seen in the results so far, a higher fraction of organic subsidies can increase NPP, in particular in terrestrial ecosystems and under scenarios of stronger consumption and more efficient nutrient conversion by animals. Besides these food web properties, we now assess which model parameters and thus which associated ecosystem processes play a role in the productivity-subsidy quality relation. We do this using both an analytical and a numerical approach.

Starting from analytical arguments, it is useful to consider a simplified scenario where predator self-regulation is negligible (i.e., *q_C_C* = 0). Not only is this approximation mathematically convenient, but this term is also indeed substantially smaller than other terms in terrestrial ecosystems (see Methods). Under this approximation, at equilibrium 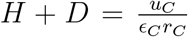, meaning that increases in *H* are balanced by decreases in *D* and vice versa as *ψ* is increased, because their total is fixed. It is therefore natural to ask how *P* changes between the low *ψ* value where *D* = 0 and the high *ψ* value where *H* = 0 (note that *D* and *H* do not necessarily span this whole range for 0 < *ψ* < 1, but this remains a useful exercise to clarify the role of some parameters). At low *ψ* and with *D* = 0, all nitrogen subsidies flow into *N*, and from eq. 1a we have at equilibrium:

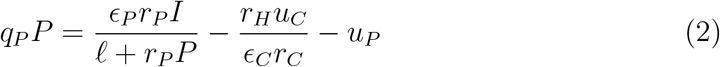

At high *ψ* and with *H* = 0, we have no herbivory term in eq. 1a, so that at equilibrium:

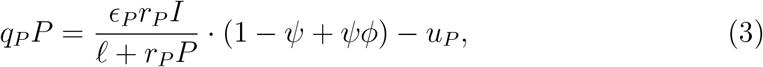

where for convenience we defined the parameter group 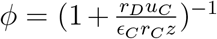. Equations 2 and 3 can be directly solved to find P, but it is more instructive to study their right hand sides, where different state variables and parameters affecting productivity *q_P_P* appear. Using these equations, we can estimate the sensitivity of productivity to changes in *ψ* because eq. 2 and 3 represent *P* at low and high *ψ*, respectively, and the derivative 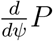 scales as *P*(*ψ* ≈ 1) – *P*(*ψ* ≈ 0).

Plant biomass *P* (and hence NPP) is lower in the low *ψ* case compared to the high *ψ* case because of the negative herbivory term in eq. 2 (i.e., 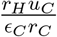). In contrast, *P* decreases in the high *ψ* case of eq. 3 when *φ* becomes smaller than 1, which reduces the term 1 – *ψ* + *ψφ*, in turn reducing the overall growth term (i.e., 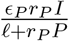). Therefore, both a larger herbivory term (which lowers *P*(*ψ* ≈ 0)) and a larger value of *φ* (which increases *P*(*ψ* ≈ 1)) lead to a more positive dependence of *P* on *ψ* (i.e., a more positive 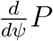).

The comparison of eq. 2 to eq. 3 thus reveals how NPP is affected by larger fractions of organic inputs, since by their construction, eq. 2 occurs for lower *ψ* values than eq. 3. For instance, fast rates of organic nitrogen dynamics (large *ℓ* and *z*) lead to a more positive effect of *ψ* on *P* (positive 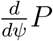). This effect occurs because larger *ℓ* decreases the overall growth term (and thereby decrease its importance relative to the herbivory term) and large *z* keeps *φ* closer to 1 (decreasing the losses due to resource competition). In both cases, less nitrogen is lost due to resource competition with detritivores, and hence NPP gains due to lower herbivory become more important. Decreasing subsidy strength *I* has the same qualitative effect as increasing *ℓ* (i.e., decreasing the relative importance of the growth term), and hence also leads to a more positive 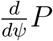.

Focusing on the herbivory term, higher herbivory rate *r_H_* also leads to a more positive 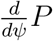, because large *r_H_* decreases *P* at low *ψ* (eq. 2). In other words, at high herbivory, *P* can increase more when eliminating herbivory by increasing *ψ* (eq. 3). Finally, higher *r_D_* decreases nutrient availability for producers by routing nutrients to decomposers (decreasing *φ* in eq. 3), thus also lowering 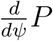. Effects of other parameters on productivity are harder to predict – for example, *u_C_* occurs in both the herbivory term and in *φ*, makings its effect less straight-foward.

### 4.4 Numerical exploration of ecosystem processes driving productivity-subsidy relations

We corroborate these analytical relationships with an extensive numerical exploration, using randomly chosen parameters sets. In this exploration, we show how the metric PSOS, which represents the average sensitivity of producer biomass *P* on subsidy quality *ψ*, varies as a function of parameter values (Fig. 4). We focus here on parameters characterizing the terrestrial ecosystem (see Table 1). A similar exploration of parameters for aquatic ecosystems shows overall comparable results (see SI), with a few exceptions, notably that in aquatic ecosystems organic additions have a more negative effect on NPP.

**Figure 4:**
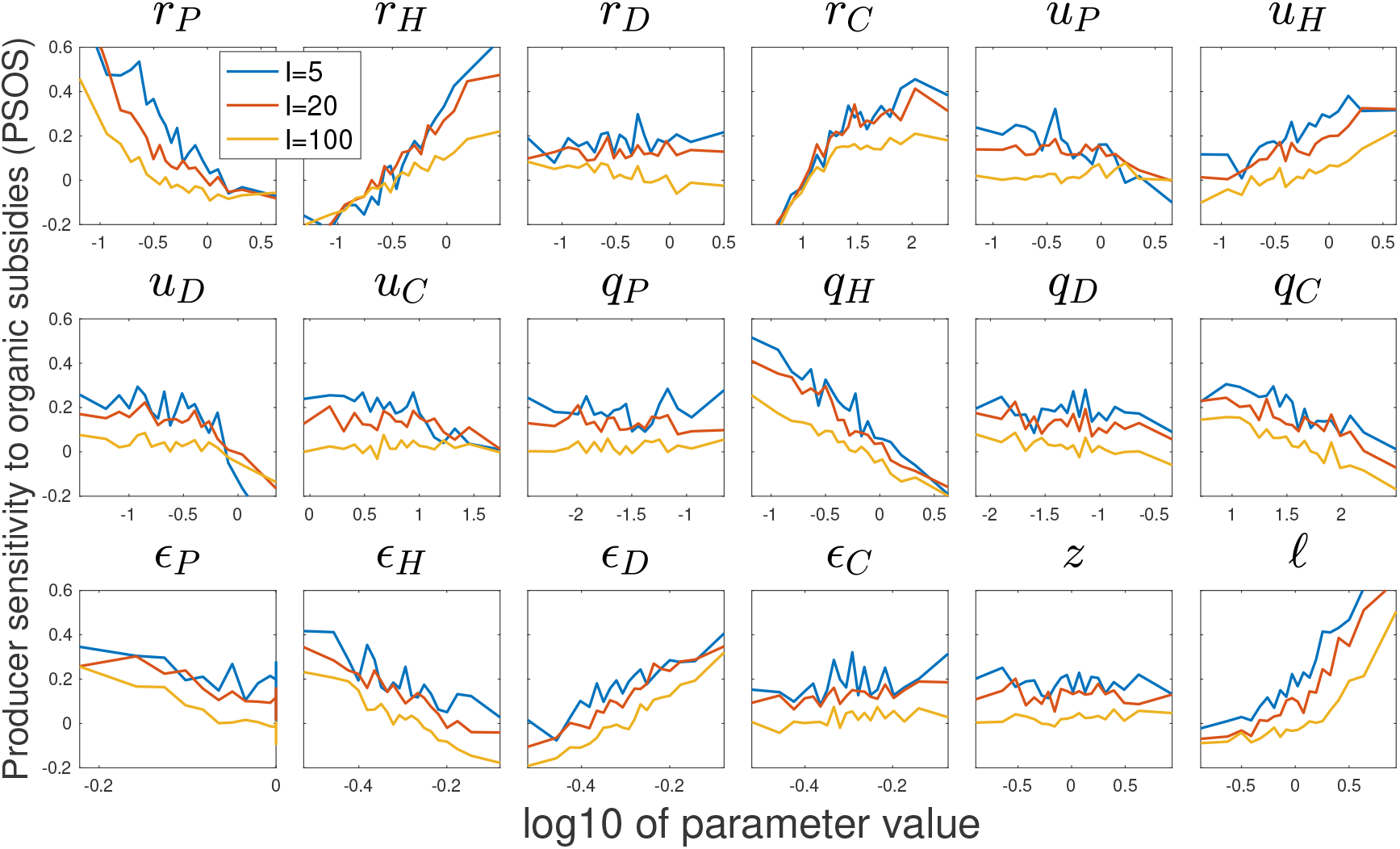
Producer sensitivity to organic subsidies (PSOS) as a function of variation in model parameters, averaged over 1000 random parameter sets per point shown. Median parameter values for the terrestrial ecosystem are used to construct the random parameter space (see Table 1). Results are shown for varying subsidy strength *I*, as indicated by different colors: blue (*I* = 5), red (*I* = 20), and yellow (*I* = 100). PSOS is defined as: 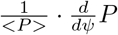.

PSOS increases with increasing *r_H_* and *ℓ*, and decreases with *I*, consistent with our analytical analysis (Fig. 4). However, *r_D_* and *z* show the predicted trends only for parameters of aquatic ecosystems (and even then with weak trends, see SI). Moreover, PSOS increases with *u_H_*, *ϵ_D_*, and *r_C_*, and decreases with *r_P_, ϵ_H_*, and *u_D_*. Additionally, self-regulation of all animal compartments causes a weak negative trend, which virtually disappears in the aquatic ecosystems (see SI).

## 5 Discussion

Our main results can be summarized along four main points. First, as expected, increasing (strengthening) nitrogen inputs leads to higher net primary productivity (NPP), but this effect saturates as nutrients inputs increase, as can be seen by the decreasing NPP values at higher subsidy strength (Fig. 2). Second, we predict that increasing the fraction of nitrogen subsidies in organic form (higher *ψ*) will lead to higher NPP in many terrestrial ecosystems, but more often to lower productivity in aquatic ecosystems (Fig. 2, Fig. S6). Third, the slope of the productivity - subsidy quality relation increases with higher consumption coefficients, but decreases substantially with higher metabolic rates (Fig. 3, Fig. S10). Fourth, we find that several ecosystem properties mediate the effect of subsidy quality on NPP (Fig. 4). For example, increasing *r_H_, r_C_*, *ϵ_D_*, and *ℓ*, consistently and strongly increases the positive effect of *ψ* on NPP.

All these relationships between subsidy and NPP depend on the ecosystem in question, and we confirm that terrestrial and aquatic ecosystems respond differently to subsidies (Nakano & Murakami, 2001; Shurin *et al*., 2006; Leroux & Loreau, 2008). The finding that in terrestrial ecosystems NPP is more positively affected by organic additions than in aquatic ecosystems may seem surprising, given that this effect is driven by trophic cascades of predators suppressing herbivores – a process that is typically associated with aquatic ecosystems. One explanation could be that metabolic rates of herbivores are often higher in aquatic ecosystems (Cebrian & Lartigue, 2004), and, as we have seen (Fig. 3), high metabolic rates lead to a more negative effect of organic inputs on NPP. We also found that increasing *z, r_P_, u_H_* and *q_H_* lead to a more positive relationship between *ψ* and NPP in aquatic ecosystems (Fig. S5). This in turn implies that in aquatic systems primary producers are nutrient limited (as increasing *z* or *r_P_* leads to stronger nutrient uptake by producers) and that herbivores are already predator-controlled (as increasing *u_H_* or *q_H_* decreases the predator control over herbivores). Hence, subsidizing predators further in aquatic ecosystems is not likely to increase NPP. Finally, we note that aquatic ecosystems often receive substantial subsidies from nearby terrestrial ecosystems (Shurin *et al*., 2006; Leroux & Loreau, 2008), and high subsidy strength *I* weakens the importance of subsidy quality *ψ* (Fig. 4). Therefore, aquatic ecosystems with high values of *I* are not likely to exhibit higher NPP due to higher fractions of organic subsidies.

Our numerical exploration (Fig. 4) largely confirms our analytical results (using eqs. 2 and 3) that low *I* and high *ℓ* lead to a positive relationship between the fraction of organic subsidies and net primary productivity. However, our analytical analysis did not allow for identifying effects of other ecosystem parameters on the relationship between subsidy quality and NPP. The numerical analysis instead pointed to the negative impact of *ϵ_H_* and *u_D_* and the positive impact of *ϵ_D_* and *u_H_*. Both high *u_H_* and low *ϵ_H_* decrease herbivore biomass, whereas low *u_D_* and high *ϵ_D_* increase detritivore biomass. All these changes lead to higher NPP when increasing organic subsidies. We thus find an asymmetry between the impacts on NPP of herbivores (which directly decrease NPP) and detritivores (which indirectly increase NPP by promoting predators).

### 5.1 Nitrogen recycling and other model assumptions

To ease the analysis and presentation, we chose not to include nitrogen recycling from producers and other compartments to the organic nitrogen compartment, which are often considered in theoretical analyses of ecosystem dynamics (De Mazancourt *et al*., 1998; Attayde & Ripa, 2008; Cherif & Loreau, 2009). However, as we show in the SI, the qualitative results are similar with and without nitrogen recycling, with three notable exceptions. Adding nitrogen recycling in the model results in higher productivity, leads to a slight shift left on the *ψ* axis as recycled material feeds the organic nutrient compartment, and weakens the positive link between *ψ* and NPP. This last aspect implies that in ecosystems where nutrient recycling is minimal, such as in agricultural crop fields, switching from inorganic to organic nutrient inputs is more likely to increase NPP.

Beyond nutrient recycling, we have made some other notable simplifying assumptions. We chose to use linear kinetics for the organic nitrogen compartment, a type *I* functional response for species interactions, and constant carbon-to-nutrient ratios in all compartments. The assumption of stoichiometric homeostasis is supported by evidence for animals, but less so for primary producers (Elser *et al*., 2000). While all of these are expected to have some effect, we expect them to make second-order corrections and they thereby should affect the quantitative results but not the qualitative ones, but this would need to be confirmed in further analyses. Since we focus here on the qualitative response of productivity to subsidies, we believe that, much like for nutrient recycling, this will not impact our general conclusions.

### 5.2 Implications and future directions

An understanding of which parts of the parameter space that are relevant to real ecosystems will be useful for applied purposes. For instance, the model predicts that introducing organic fertilization in agriculture improves regulation of herbivores, which are potential pests on crop plants. Further, in conjunction with site-specific parameter estimates, the model we present can be used to predict the effects of anthropogenic subsidies on specific natural ecosystems, e.g., grasslands and forests, as it would clarify which ecosystems are more sensitive to nutrient enrichment. Finally, climate change affects metabolic rates and the kinetics of soil nutrient cycling, which in turn affect the impacts of nutrient subsidies on NPP. This highlights the importance of exploring how ecosystems respond to compound climatic changes and perturbations in the subsidy strength and quality. These changes lead ecosystems towards conditions not previous experienced, which are therefore also harder to predict. Our approach, of developing theory for subsidized ecosystems using dynamical models, parameterizing these models for a wide range of ecosystem, and testing confounding effects of model parameters, is therefore a crucial step in understanding how ecosystems respond to human influence. Importantly, it is crucial to consider the green and brown food webs in concert.

While useful for qualitative predictions, testing our theoretical predictions empirically in specific case studies is likely to be challenging. There are recent investigations of nutrient quality and quantity subsidy effects on primary producers in conjunction with food-web dynamics (Riggi & Bommarco, 2019; Aguilera *et al*., 2021), but it is difficult to directly connect their results to our theoretical findings. A particular problem is to normalize results by subsidy strength, i.e., empirically disentangling *I* and *ψ*. This is of interest since the subsidy effect in the context of green-brown good webs should be most striking along the *ψ* axis, given that the effect of *I* on productivity is a more straightforward bottom-up one. Indeed, despite clear results showing that detritus additions increase primary producer stocks (Hagen *et al*., 2012), it is not possible to attribute subsidy effects to bottom-up vs. top-down mechanisms, because experiments typically manipulate both *ψ* and *I*, and increases in *I* may overshadow the top-down effects of higher *ψ*. A partial solution is to compare ecosystems with and without herbivory and/or predation (e.g., due to pesticide use), where the effect of *ψ* for different parameter regimes should be most evident.

Here we explored how two aspects of nutrient subsidies – strength and quality – on primary production in green-brown food webs. Subsidy strength has a direct positive impact, mainly as a bottom-up direct effect on producers. The effect of subsidy quality truly connects the green and brown channels in food webs – organic subsidies can indirectly promote productivity via predator control on herbivores. This aspect of nutrient enrichment has been largely overlooked, and as we have seen here can play a major role in determining ecosystem functioning. These results show that the impact of nutrient enrichment depends on more than its strength, and that human overloading of ecosystems with inorganic nutrients is consequential not only because of its large amounts, but also due to higher proportions of inorganic nutrients, which promote productivity at the cost of losing the ecological function of the brown channel.

## 5.3 Acknowledgements

This research was funded by the Swedish University of Agricultural Sciences, and by the European Research Council (ERC) under the European Union?s Horizon 2020 research and innovation programme (grant agreement No 101001608).

## Supporting Information

In this Supporting Information, we complement and generalize the results shown in the main text. First, the assumption of negligible nutrient recycling is relaxed; second, results shown in Figs. 3-4 for terrestrial ecosystems are complemented by analyses for aquatic ecosystems (Table 1); third, the ramification of using a type-II functional response are tested; fourth, uncertainties in the results due to the parameterization of consumption, conversion and metabolic rates are assessed.

### Nutrient recycling and aquatic ecosystems

We test the impact of including explicit nutrient recycling to the model, by adding nutrient recycling from primary producer mortality. We do this by changing the value of the recycling fraction *y* in the input term *I*_0_ of eq. 1f, where *I*_0_ = *yu_P_P*. We calculate the nitrogen recycling fraction as *y* = 1 – *f_EX_/f_P_*, where *f_EX_* represents how much biomass is exported out of the ecosystem and *F_P_* represents the primary production flux. Both fluxes are obtained from Cebrian dataset (see Methods). This calculation gives *y* = 0.982 for the terrestrial ecosystem and *y* = 0.815 for the aquatic one. Next, the effect of nitrogen recycling is assessed by reproducing Fig. 2 and Fig. 4, but now including the new recycling term. In Fig. S1, the effect of subsidies in primary production are shown, using the parameters for both terrestrial and aquatic ecosystems.

In comparison to Fig. 2, all the qualitative behaviors are the same. The two main differences are that overall production levels are higher (since the ecosystem is more efficient in retaining and reusing nitrogen) and that there is a shift left along the subsidy quality axis (*ψ*). This shift is most clearly discernible by looking at how the coexistence range, marked by gray lines, has moved to the left. The leftward shift is due to the recycled material entering the organic nitrogen compartment, which becomes less dependent on external organic subsidies. As a result, when recycling occurs, the simulated ecosystem reaches at low organic subsidy (i.e., low *ψ*) a similar state as with intermediate organic subsidy, but without recycling.

Next, we extend the results of Fig. 4, in which we considered the effect of organic subsidies on primary productivity, confounded by different model parameters, using the metric Producer Sensitivity to Organic Subsidy (PSOS). In Fig. S2 we reproduce the numerical exploration of Fig. 4, but now with the recycling term. We also conduct this numerical exploration for random parameter sets centered on the parameters for the aquatic ecosystem (Fig. S3), as well as when adding nutrient recycling to the aquatic ecosystem (Fig. S4). In all three cases the results are quite similar to those shown in the main text.

A summary of all ecosystem types (corresponding to Fig. S2, Fig. S3, and Fig. S4, as well as Fig. 4 in the main text), that compares these different parameterization options (aquatic/terrestrial; with/without nutrient recylcing), is given in Fig. S5 and Fig. S6. In Fig. S5 we show how PSOS depends on each one of the 18 parameters that were varied. For each parameter, we calculate a linear fitting for PSOS as a function of log10 of the parameter value, for a given value of *I*. The slopes of these linear fits are calculated for each of the three values of *I* used in the aforementioned figures, namely 5, 20, and 100 [*gNm*^−2^*yr*^−1^]. The average of the three slopes is then used in Fig. S5. In Fig. S6 we show the average value of PSOS for different values of *I* (1, 2, 5, 10, 20, 50, 100 [*gNm*^−2^*yr*^−1^]). For each of the four ecosystem types (terrestrial/aquatic, with/without nutrient recycling), we average over the PSOS values corresponding to the 20000 different parameter sets for each ecosystem type.

**Figure S1:**
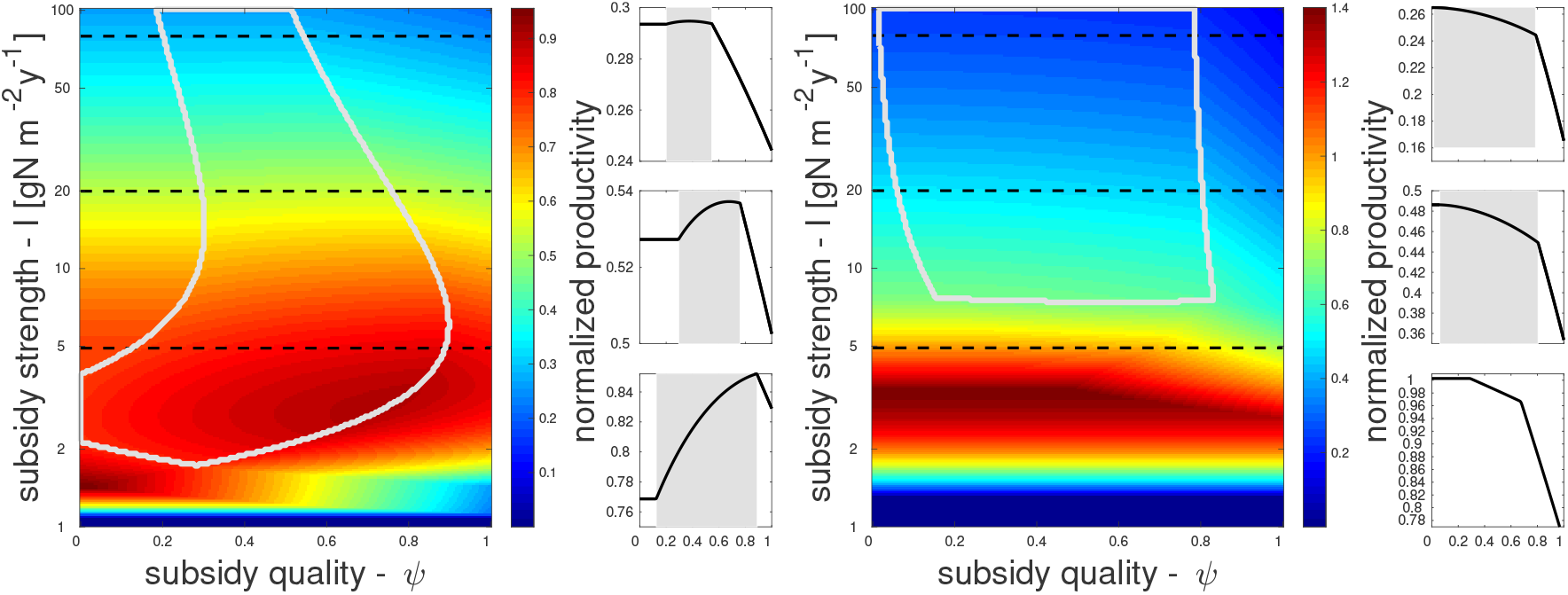
Normalized primary production *u_P_P/I* as a function of nutrient subsidies, *I* (strength) and *ψ* (organic fraction), for the model including explicit recycling of dead primary producer material. Besides *I* and *ψ*, parameters are constant, corresponding to the terrestrial (left) and aquatic (right) ecosystems. Gray lines show the coexistence region where all food web compartments (*P,H,D,C*) are extant. Small panels on the right of each parameter-space show primary productivity at a constant value of *I* (gray background shows coexistence region), and corresponding to the three horizontal dashed lines in each parameter-space. Fraction of dead primary producer material flux that is recycled is *y* = 0.982 (*y* = 0.815) for the terrestrial (aquatic) ecosystem.

**Figure S2:**
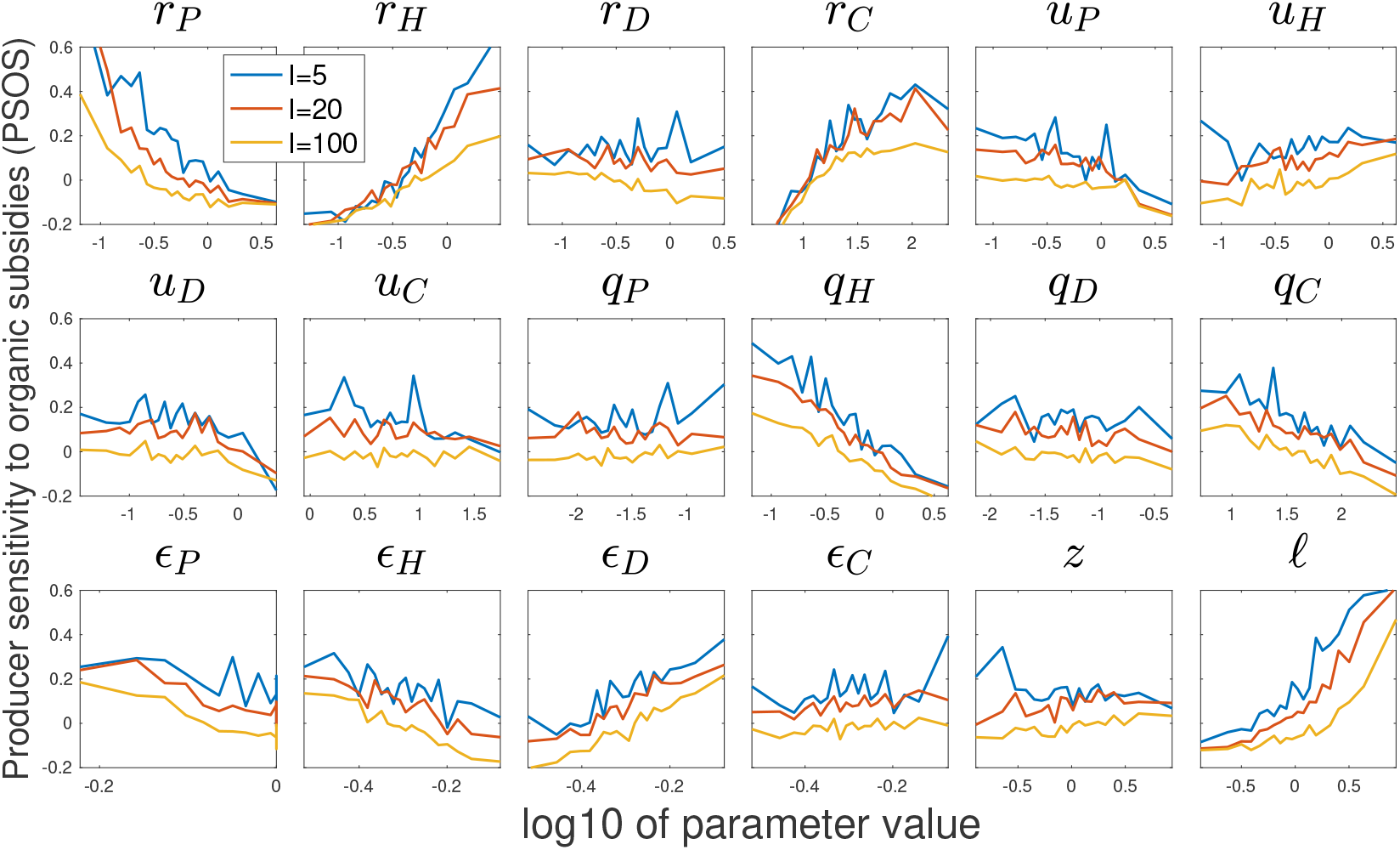
Producer sensitivity to organic subsidies (PSOS) for terrestrial ecosystems with explicit nutrient recycling. Averaged PSOS is shown as a function of different model parameters, averaged over 1000 random parameter sets per point shown. Results are shown for increasing subsidy strength *I* with different colors: blue (*I* = 5), red (*I* = 20), and yellow (*I* = 100) [*gNm^−2^*yr*^−1^*].

**Figure S3:**
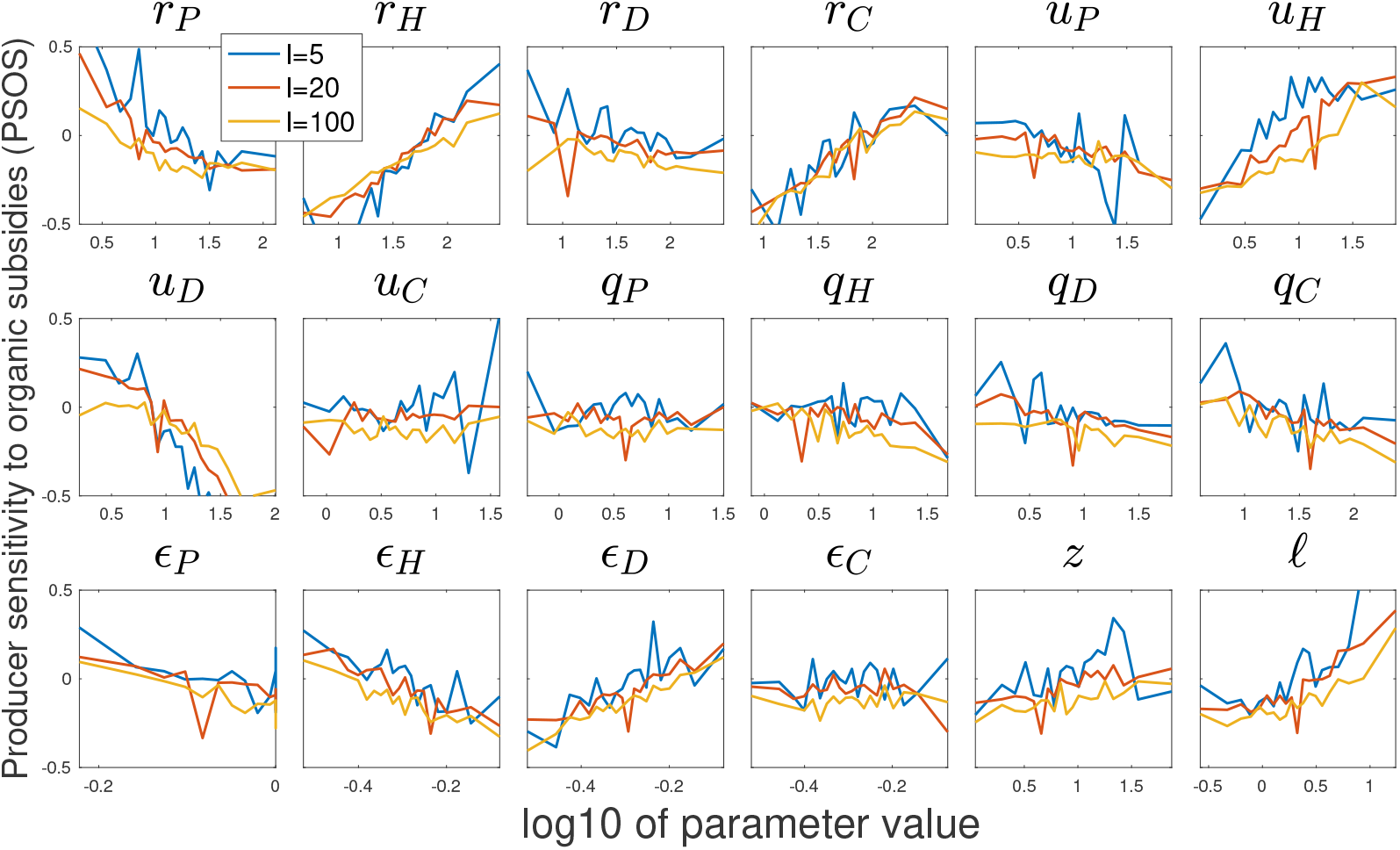
Producer sensitivity to organic subsidies (PSOS) for aquatic ecosystems without nutrient recycling (see Table 1). Average PSOS is shown as a function of different model parameters, averaged over 1000 random parameter sets per point shown. Results are shown for increasing subsidy strength *I* with different colors: blue (*I* = 5), red (*I* = 20), and yellow (*I* = 100) [*gNm*^−2^*yr*^−1^]

**Figure S4:**
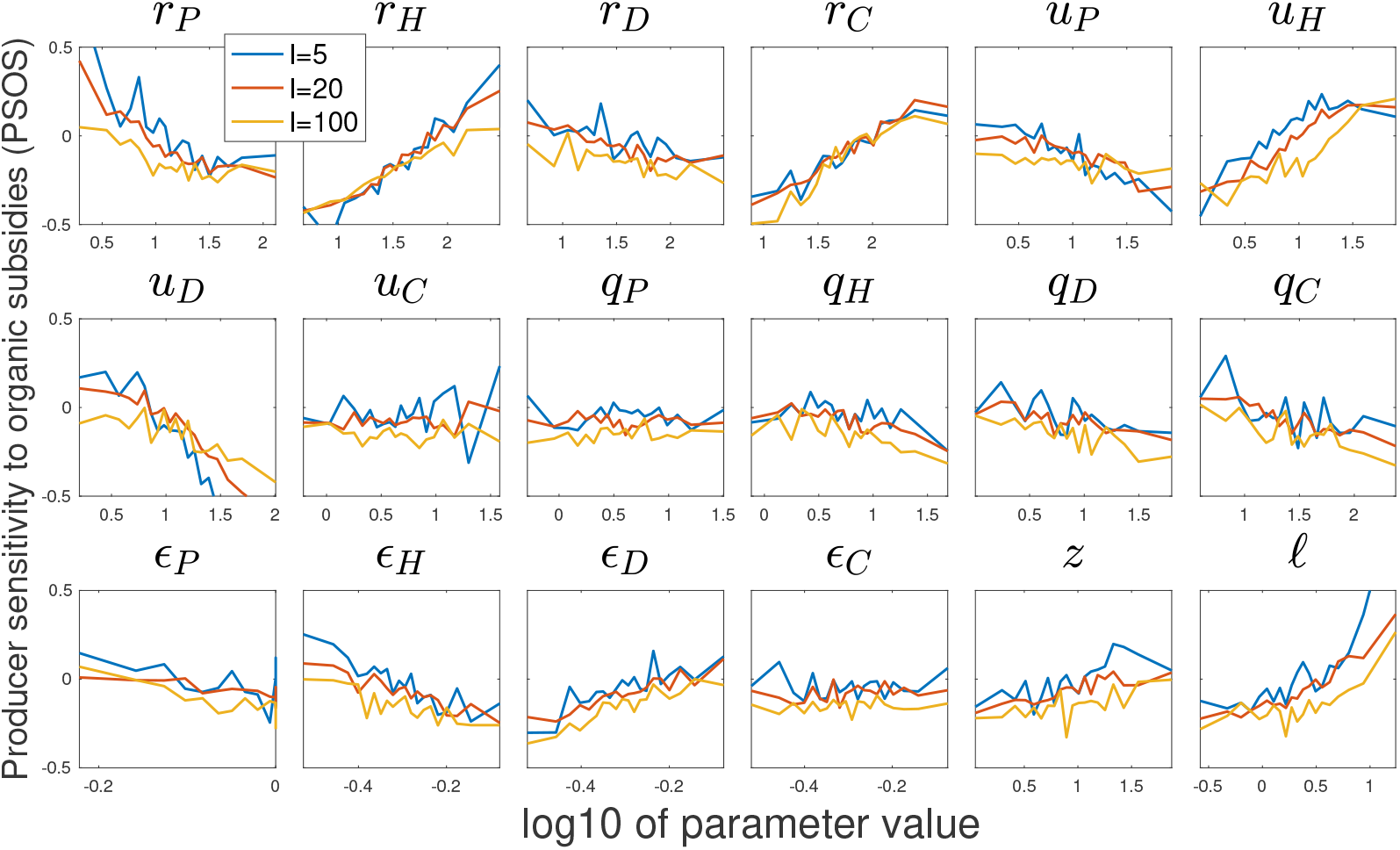
Producer sensitivity to organic subsidies (PSOS) for aquatic ecosystems with explicit nutrient recycling (see Table 1). Averaged PSOS is shown as a function of different model parameters, averaged over 1000 random parameter sets per point shown. Results are shown for increasing subsidy strength *I* with different colors: blue (*I* = 5), red (*I* = 20), and yellow (*I* = 100) [*gNm*^−2^*yr*^−1^]

Notably, the effect of recycling on parameter dependency (closed versus open circles in Fig. S5) is small compared to the overall variability among different parameters and to the differences between terrestrial and aquatic ecosystems. The average values of PSOS are noticeably lower when recycling is included, and they decrease as subsidy strength *I* increases (Fig. S6). This shows that in ecosystems with less recycling, the positive effect of organic additions on primary production is expected to be considerably stronger. However, at very high subsidy strength, the importance of recycling decreases.

From these analyses, several differences emerge between terrestrial and aquatic ecosystems. Most notably, the average values of PSOS are positive for terrestrial but mostly negative for aquatic ecosystems (Fig. S6). Several parameter dependencies (Fig. S5) are stronger in aquatic ecosystems – *u_H_* and *u_D_*, and less significantly for *r_D_* and *z*. The opposite trend of weaker dependency for aquatic ecosystems is seen for *r_P_*, *q_H_*, and *ℓ*.

**Figure S5:**
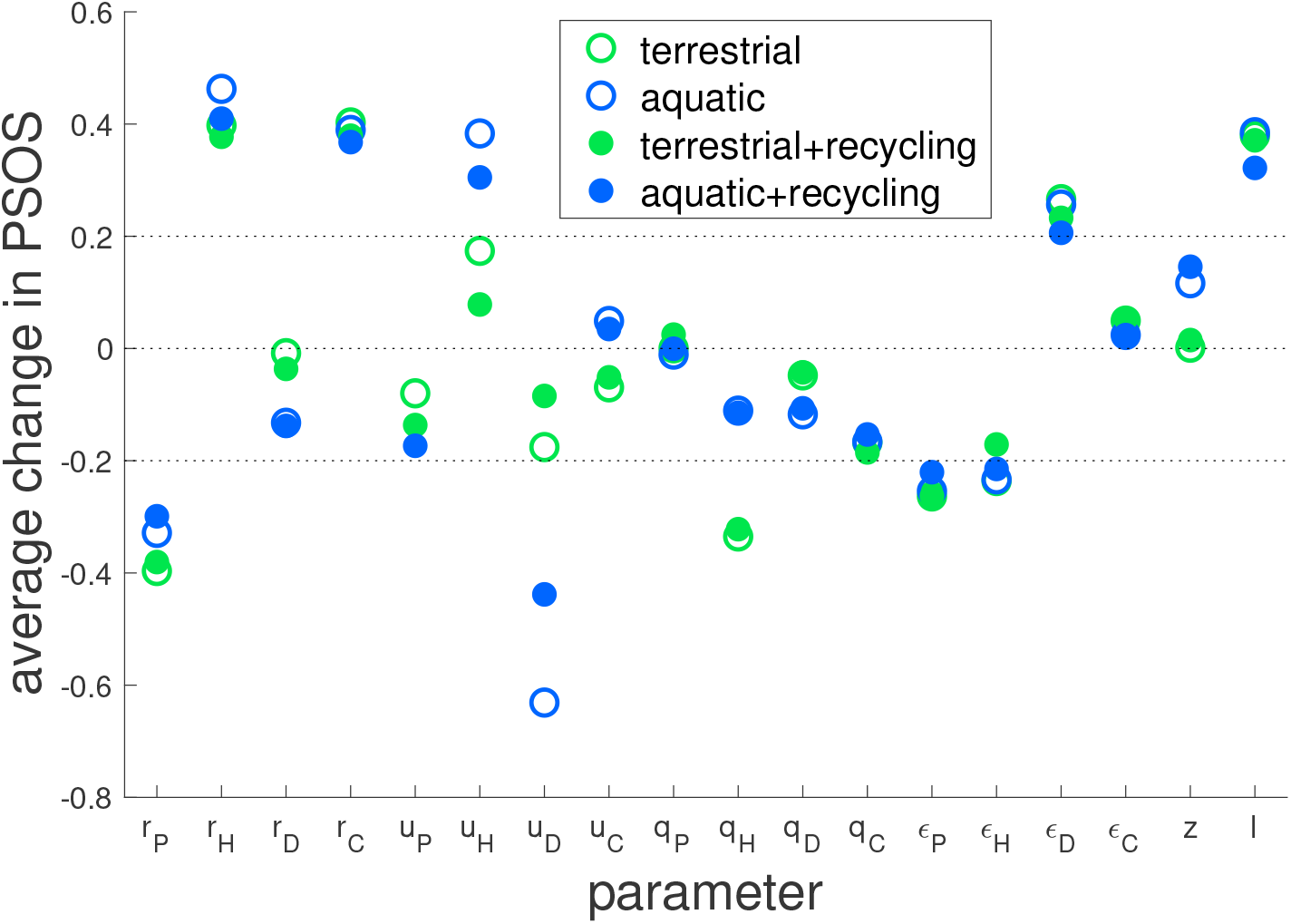
Average change in producer sensitivity to organic subsidies (PSOS) when increasing a model parameter, for different model parameters. Average change is shown for four types of ecosystems, aquatic (blue) and terrestrial (green), with and without explicit primary producer nutrient recycling (closed and open circles, respectively). For each ecosystem type, one of 18 parameters, and constant value of subsidy strength, the average change is calculated as the slope for a linear fit of PSOS as a function of log10 of the given parameter. Values shown are the averages over three values of subsidy strength: *I* = 5, *I* = 20, and *I* = 100 [*gNm*^−2^*yr*^−1^].

**Figure S6:**
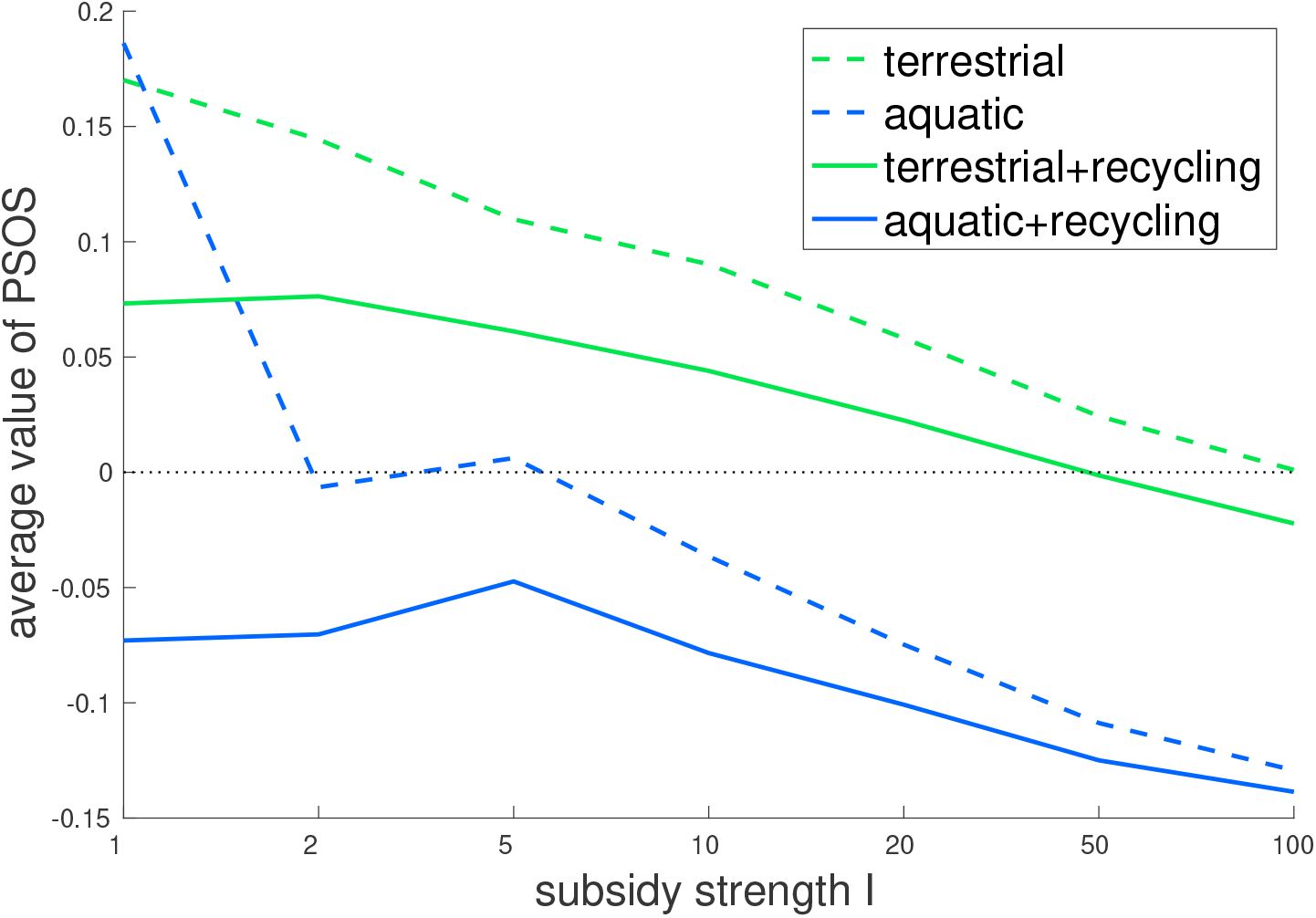
Average producer sensitivity to organic subsidies (PSOS) as a function of subsidy strength. Average PSOS is shown for four types of ecosystems, aquatic (blue) and terrestrial (green), with and without explicit primary producer nutrient recycling (dashed and solid lines, respectively). For each ecosystem type and value of *I*, PSOS values are averaged over all parameter sets where PSOS can be calculated (in particular, where all four food web compartments are extant).

### Comparing type-I to type-II functional response

We compare here our model in the main text of a type-I functional response for consumption, to an alternative model of a type-II functional response.

In the results shown in the main text, we use a type-I functional response for consumption, e.g., the term *P* · *ϵ_P_r_P_N* in eq. 1. We extend this model to a type-II functional response using the normalized handling time *h_i_*. For instance, for *P* we do this by dividing the original consumption term by a term of the form 1 + *h_P_N*, where *h_P_* is the normalized handling time for consumption by *P*. We thus arrive at eq. S1, where we opted to use two separate terms for the consumption by predators (*ϵ_C_r_C_CH*/(1 + *h_C_H*) and *ϵ_C_r_C_DC*/(1 + *h_C_D*)), for simplicity.

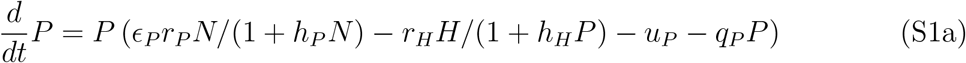

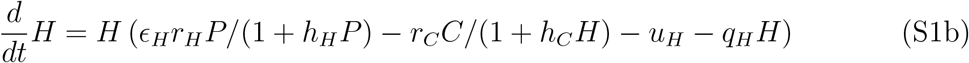

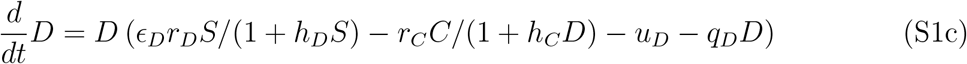

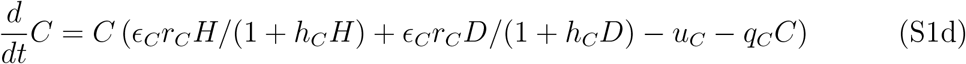

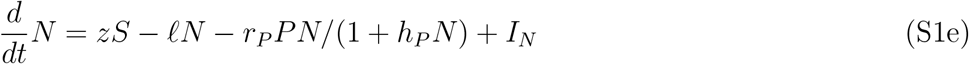

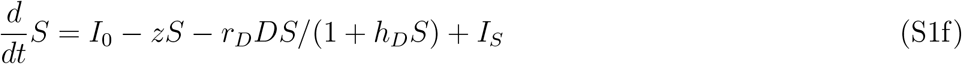

Estimating the normalized handling time *h_i_* for an entire compartment (e.g., for the whole herbivore compartment, and not for a specific species of herbivores) is not straight forward, and we therefore define the values for *h_i_* based on two considerations: 1) For consistency with our model for type-I functional response, we define *h_i_* so that at ambient values, i.e., the stocks and fluxes values at which we calculate parameters, the consumption flux is identical for both type I and type II. 2) The degree on non-linearity of the type-II response is defined by *n*: how much larger the per-capita maximal consumption flux is relative to ambient levels, where a smaller ratio *n* is associated with a stronger non-linearity.

Given these two considerations, we arrive at 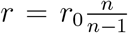 and 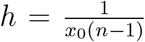. Here *x*_0_ is the stock of the resource at ambient levels, r_0_ is the value for the consumption coefficient for the type-I functional response, and *r* and *h* are the values used for the consumption coefficient and the normalized handling time in the new model, respectively. For instance, with *n* = 2 and a given level of consumer stocks, the saturation of consumption flux with higher resource stocks is strong, and the flux can at most double, relative to ambient levels.

We simulate the type-II model similarly to the main text, with a set of 20000 simulations such as for Fig. 4, for both weak (*n* = 4) and strong (*n* = 2) nonlinearity of the functional response. For each set, we define the median values of the normalized handling time parameters *h_i_* using the definition above and either *n* = 4 or *n* = 2. We then proceed to randomly choose the values of *h_i_* in the same manner as for the other parameters, by choosing a random log-normal distribution around the median value, such that the base-10 logarithm values have a standard-deviation of 0.5.

We test the validity of these simulations in two ways, by checking first if all compartments are extant, i.e., that the stocks of *P, H, D* and *C* are all above zero, and by checking if the simulations do not lead to oscillations. We note that both these conditions, no extinctions and no oscillations, are expected to hold in the context of functional groups, unlike the case of specific species, where extinctions and oscillations are more likely to occur. We consider a compartment to be extinct if its stocks is below a threshold of 10^−4^ (where typical non-extinct values are of the order of 1), and we consider the system to be oscillating if the simulations has not reached an equilibrium after 1000 years (where most simulations reach an equilibrium in under 100 years).

We find that with a type-II response, a much smaller fraction of simulations end with extant communities (Fig. S7), in particular for parameters corresponding to the terrestrial ecosystem. Further, unrealistic oscillations are between 4 and 15 times more common with a type-II response (Table S1).

Overall, given that a type-II functional response is more likely to lead to compartment extinctions and to oscillations when compared with a type-I functional response, we conclude that a type-I is more appropriate for our ecosystem model.

**Figure S7:**
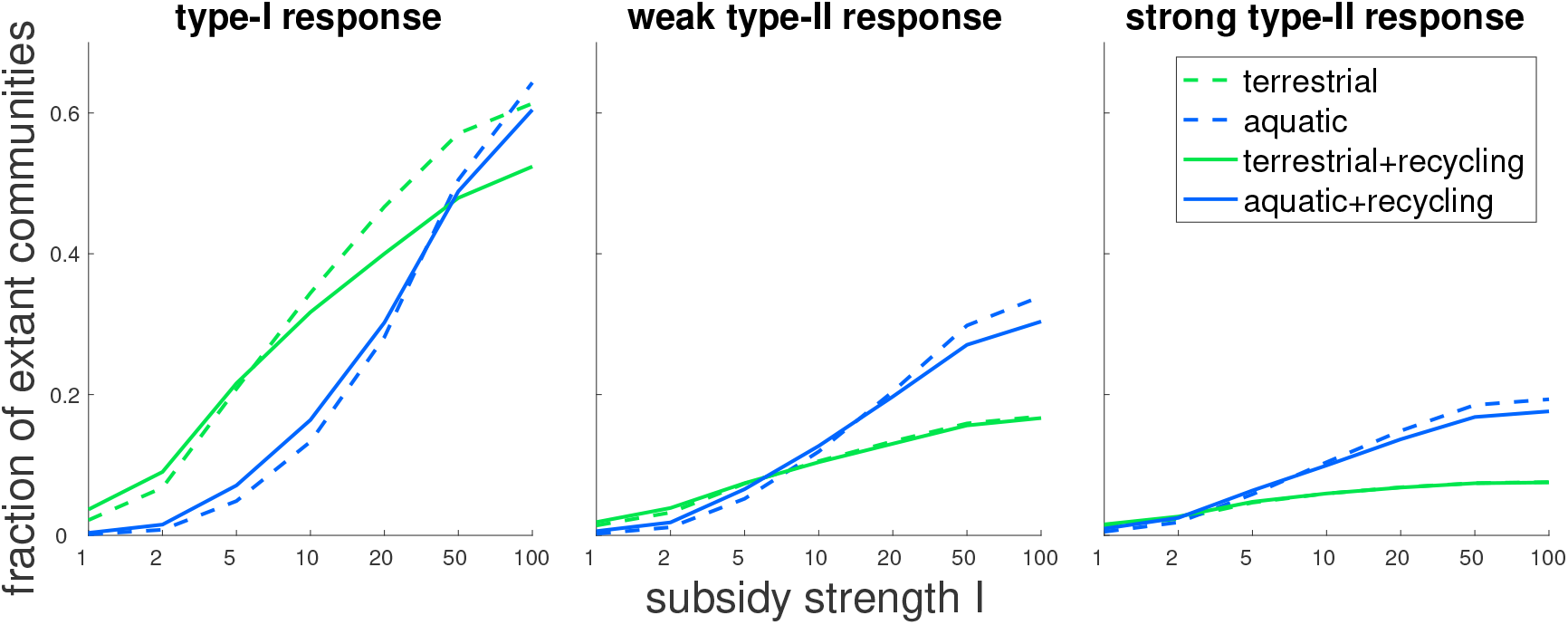
The fraction of simulations resulting in a fully extant community, with the stocks of *P*, *H*, *D* and *C* all above zero. Green (blue) lines denote parameters for terrestrial (aquatic) ecosystems, and dashed (solid) lines denote models without (with) nutrient recycling. Left panel is for the model in the main text with a type-I functional response, while middle and left panels are for a type-II functional response, with a weak (*n* = 4) and strong (*n* = 2) nonlinearity, respectively.

**Table S1:** Percent of simulations resulting in temporal oscillations. Rows correspond to parameters for either terrestrial or aquatic ecosystems, with or without nutrient recycling. Left column is for the model in the main text with a type-I functional response, while middle and left columns are for a type-II functional response, with a weak (*n* = 4) and strong (*n* = 2) non-linearity, respectively.

| | type-I | weak type-II ($n=4$) | strong type-II ($n=2$) |
| --- | --- | --- | --- |
| terrestrial | 0.7 | 5.5 | 4.2 |
| aquatic | 1.1 | 15.6 | 14.8 |
| terrestrial + recycling | 0.9 | 4.8 | 3.9 |
| aquatic + recycling | 1.1 | 14.7 | 13.5 |

### Effect of food web properties

In this section, we test if our choice for the parameterization of food web properties alters the trends found in Fig. 3. To this aim, additional parameterization schemes for consumption, metabolic rates, and conversion are defined (Table S2); moreover, alternative baseline parameter values are considered for the ecosystems tested.

In each of the nine parameterization schemes shown in Table S2 (the three used in main text, and six additional ones tested here), we rescale several different model parameters, using a scaling factor *γ*. Beyond the original three schemes (noted as c1, m1 and a1 in Table S2), we use six other schemes, by: 1) also rescaling the parameters for primary producers (and not only for animal consumers) – in c2, c4, m3, and a2; 2) defining the food web property of consumption by rescaling self-regulation in addition to consumption coefficients – in c3 and c4; 3) increasing metabolic rates differently for predators – in m2. All nine parameterization schemes are presented in Table S2. Note that for the additional parameterization of conversion (a2) we first re-scale the baseline value of *ϵ_P_* to 0.7, and we also halve its scaling constant, in order to limit its range to high values close to 1, but not above it. We compare between these parameterization schemes in Fig. S8 (for subsidy quality *ψ*) and Fig. S9 (for subsidy strength I). In these figures the results for (c1, m1, a1) correspond to the results shown in Fig. 3 in the main text.

**Table S2:** Factors applied on specific parameter values, for the nine food web properties tested. Consumption coefficients: c1-c4, metabolic rates: m1-m3, nutrient conversion efficiencies: a1-a2. As noted in the Methods, the values of *γ* are: 0.1 (very low), 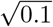 (low), 1 (medium), 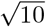 (high), and 10 (very high). Food web properties in bold (c1, m1, a1) correspond to the results shown in the main text (Fig. 3).

|  | Food web property |  |  |  |  |  |  |  |  |
| --- | --- | --- | --- | --- | --- | --- | --- | --- | --- |
|  | consumption |  |  |  | metabolic rate |  |  | conversion |  |
| parameter | **c1** | c2 | c3 | c4 | **m1** | m2 | m3 | **a1** | a2 |
| $r_P$ | | $\gamma$ | | $\gamma^{1/2}$ | | | $\gamma$ | | |
| $r_H$ | $\gamma$ | $\gamma$ | $\gamma^{1/2}$ | $\gamma^{1/2}$ | $\gamma$ | $\gamma$ | $\gamma$ | | |
| $r_D$ | $\gamma$ | $\gamma$ | $\gamma^{1/2}$ | $\gamma^{1/2}$ | $\gamma$ | $\gamma$ | $\gamma$ | | |
| $r_C$ | $\gamma$ | $\gamma$ | $\gamma^{1/2}$ | $\gamma^{1/2}$ | $\gamma$ | $\gamma^2$ | $\gamma$ | | |
| $u_P$ | | | | | | | $\gamma$ | | |
| $u_H$ | | | | | $\gamma$ | $\gamma$ | $\gamma$ | | |
| $u_D$ | | | | | $\gamma$ | $\gamma$ | $\gamma$ | | |
| $u_C$ | | | | | $\gamma$ | $\gamma^2$ | $\gamma$ | | |
| $q_P$ | | | | $\gamma^{-1/2}$ | | | $\gamma$ | | |
| $q_H$ | | | $\gamma^{-1/2}$ | $\gamma^{-1/2}$ | $\gamma$ | $\gamma$ | $\gamma$ | | |
| $q_D$ | | | $\gamma^{-1/2}$ | $\gamma^{-1/2}$ | $\gamma$ | $\gamma$ | $\gamma$ | | |
| $q_C$ | | | $\gamma^{-1/2}$ | $\gamma^{-1/2}$ | $\gamma$ | $\gamma^2$ | $\gamma$ | | |
| $\epsilon_P$ | | | | | | | | | $\gamma^{1/8}$ |
| $\epsilon_H$ | | | | | | | | $\gamma^{1/4}$ | $\gamma^{1/4}$ |
| $\epsilon_D$ | | | | | | | | $\gamma^{1/4}$ | $\gamma^{1/4}$ |
| $\epsilon_C$ | | | | | | | | $\gamma^{1/4}$ | $\gamma^{1/4}$ |

**Figure S8:**
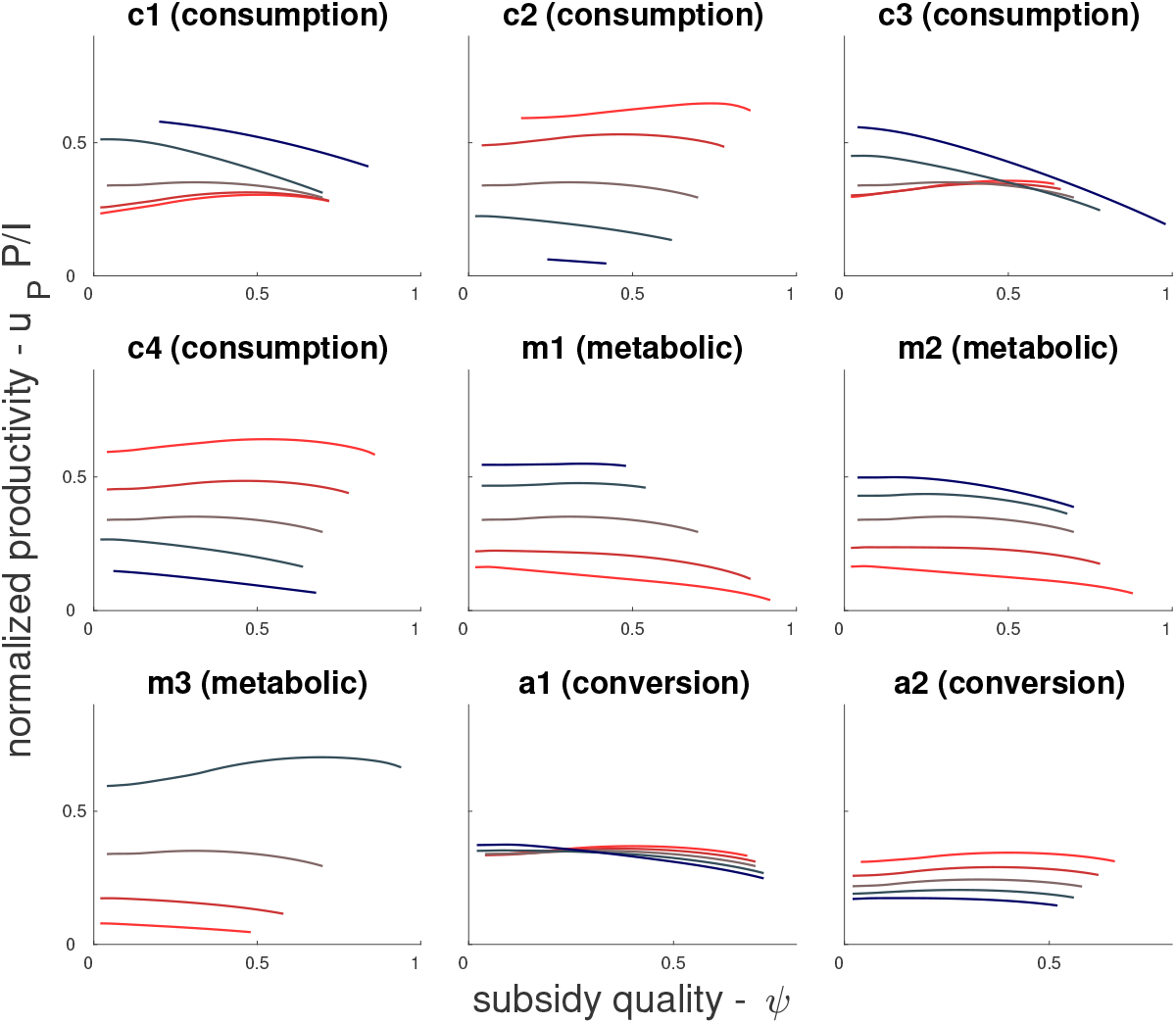
The effect of food web properties and subsidy quality on primary production. This figure shows the same results as top panels in Fig. 3, but including additional options of parameterizing consumption, conversion, and metabolic rates (as defined in Table S2).

**Figure S9:**
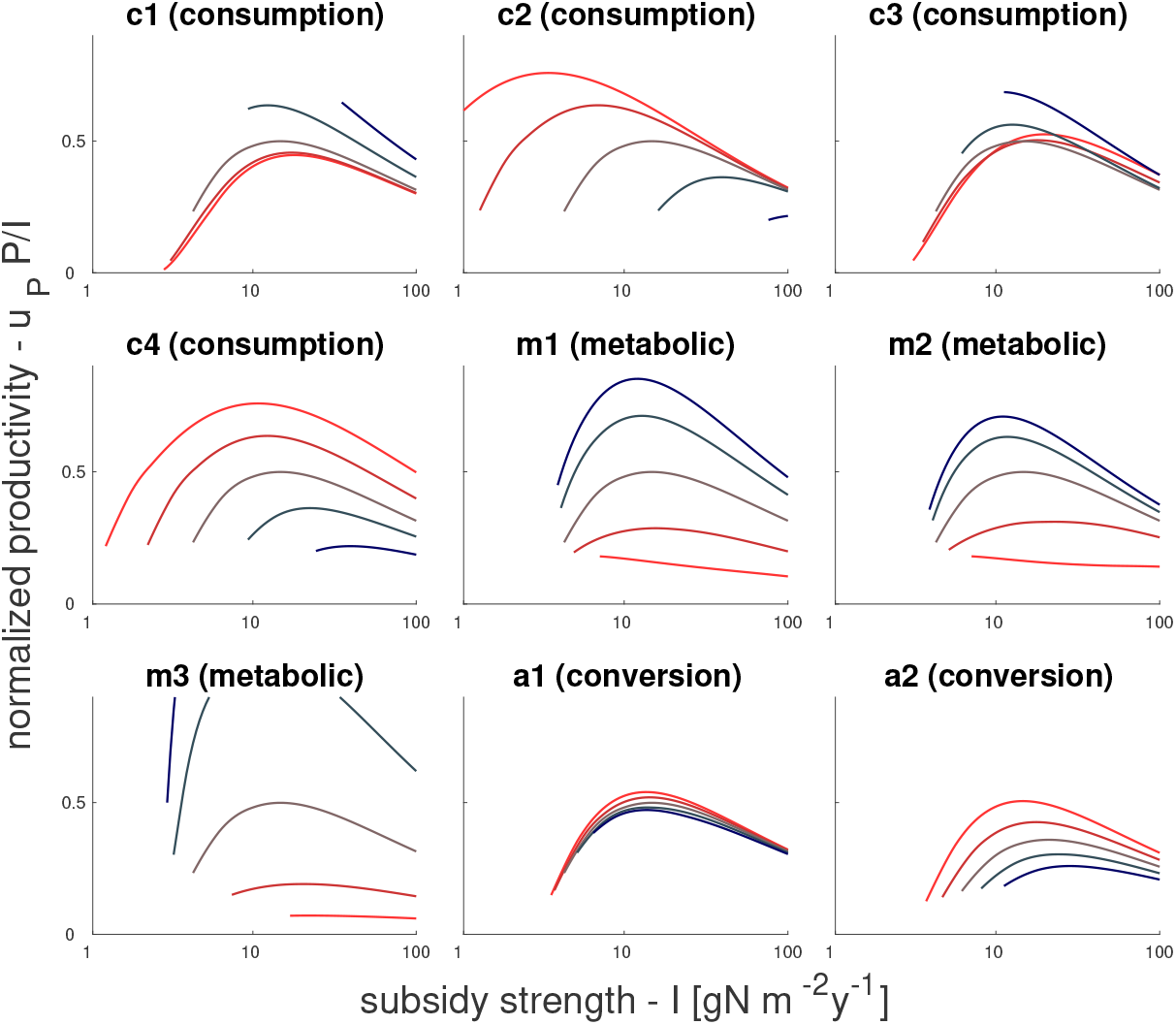
The effect of food web properties and subsidy strength on primary production. This figure shows the same results as bottom panels in Fig. 3, but including additional options of parameterizing consumption, conversion, and metabolic rates (as defined in Table S2).

We also test all nine parameterization schemes (corresponding to different food web properties) for different baseline parameters values, corresponding to different ecosystem types. We do this by assembling 160 different parameter sets that will serve as the baseline values, and test several dependencies of *P* on the nine parameterization schemes. The assembled parameters sets are composed of four groups, each having 40 parameter sets. The four groups correspond to the four main types of parameter sets explored (e.g., in Fig. S5): terrestrial ecosystems as in the main text, terrestrial ecosystems with explicit nutrient recycling, and aquatic ecosystems either with or without nutrient recycling.

For each of these four, we start with a large set of random parameter values – the 20000 random parameter sets used for the parameter exploration. We separate each such large set into two subsets, for either positive or negative PSOS, which is calculated at the medium subsidy strength of *I* =10 [*gNm*^−2^*yr*^−1^]. We are thus left with 8 subsets (four ecosystem types, and either positive or negative PSOS). In this way we make sure our results do not depend on the type of PSOS dependency.

From each of these subsets we choose 20 parameter sets, according to two criteria: 1) how different the specific parameter set is from the median parameter set (taken as the normalized distance of all 18 parameters); 2) how large the range of coexistence is along the *ψ* axis, for the medium subsidy strength of *I* =10 [*gNm*^−2^*yr*^−1^]. We choose only parameter sets for which in at least half of the *ψ* range (from 0 to 1) all compartments are extant (as explained in the main text, we can assume that realistic ecosystems have all compartments). From the parameter sets with a large coexistence range we choose the 20 parameter sets that are most similar to the median parameter set, i.e., that minimize criterion 1. In this way we make sure that the ecosystems parameter sets chosen are not extreme in their values.

With the 160 (20 parameters by 8 subsets) parameter sets and nine parameterizations, we test three aspects of productivity responses to subsidies: 1) how increasing *γ* itself changes *P* (e.g., how does increasing metabolic rates affect P); 2) how does changing *γ* affect the dependency of *P* on *ψ* (similarly to PSOS); 3) how does changing *γ* affect the dependency of *P* on *I*. For all of these, we use simulations with 5 values of *γ*, and a range of values for *I* and *ψ* (between 1 and 100 [*gNm*^−2^*yr*^−1^], and between 0 and 1, respectively). For (1), we simply average over *P* for the whole *I-ψ* parameter space, and see if the average *P* increases or decreases when we increase *γ*. For (2), we average over the *I* axis for each value of *ψ*, and consider only *ψ* values for which there is coexistence (value of all compartments is above 10^−5^[*gNm*^−2^]) for set value of *I*. We then calculate a linear fitting for *P* as a function of *ψ*, and ask if the slopes increase or decrease when increasing *γ*. For (3) we follow a similar procedure, except that we average over the *ψ* axis and evaluate slopes of fitted lines along the *I* axis.

Fig. S10 shows, for the nine different paramermization schemes, how *P* is affected by increasing the control parameter *γ*. The left and middle panels show the probability of a more positive slope of *P* as a function of *ψ* (left panel) and *I* (middle panel), while the right panel shows the probability that the increase in *γ* increases *P* directly. The results of Fig. S10 demonstrate that the following trends (which were already reported in the main text) also hold for a wider range of baseline parameter sets, representing very different ecosystems:

1. Both higher consumption and higher conversion lead to a more positive relationship between *ψ* and *P* – i.e., a larger value of PSOS. This is seen in the left panel of Fig. S10, where average probabilities of increasing slopes are between 0.7 and 0.9.
2. Higher metabolic rates lead to lower PSOS. This is not a strong effect, seen by values for (m1, m2, m3) in the left panel being only slightly lower than the neutral probability of 0.5, and is consistent with results in the main text.
3. Higher metabolic rates lead to weaker dependence on subsidy strength *I*. This is seen in the middle panel where the probabilities of increasing slope are high consistently. The high probabilities here signify a transition that occurs for medium to high values of *I* (see Fig. S9), where a very negative slope (black line) changes to an almost zero slope (red line). The same is true to a lesser degree for consumption and conversion, with the exception of a2.
4. Higher metabolic rates lower P. This result is shown in the right panel, where for m1, m2, and m3, the probabilities of increasing slope are all zero. In addition, the probabilities for c1 (the chosen approach to change the food web’s consumption rates in the main text) are low. In contrast, other parameterization options for consumption, and all options for conversion lead to higher probabilities of increasing *P* values.

**Figure S10:**
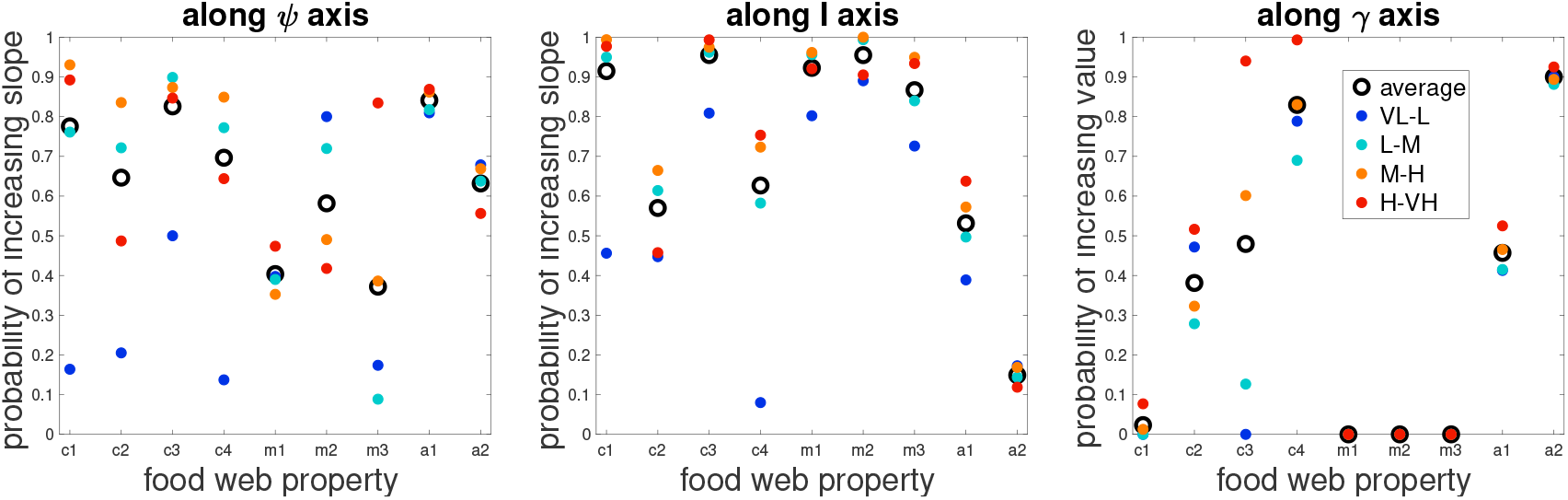
Change in primary production due to change in food web properties. In all panels, we show how changing food web properties affects *P* values, via changes in subsidy properties *ψ* and *I*, or directly. Nine different rescaling schemes for food web properties are tested, as detailed in Table S2. Different panels show probability of change in *P* as *γ* is increased (corresponding to higher food web properties), in different ways; **left**: more positive slope of *P* as a function of *ψ* (i.e., higher PSOS); **middle**: more positive slope of *P* as a function of I; **right**: higher value of P. Circles in different colors correspond to changing *γ* between five different levels: very low (VL), low (L), medium (M), high (H), very high (VH). **blue (VL-L)**: from *γ* = 0.1 to 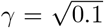; **cyan (L-M)**: from 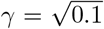 to *γ* = 1; **orange (M-H)**: from *γ* = 1 to 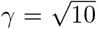; **red (H-VH)**: from 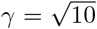 to *γ* = 10. Black rings average over all four options. For all panels and food web properties, the changes in *γ* are tested over 160 different parameter sets, corresponding to a wide array of ecosystem types, including both aquatic and terrestrial ecosystems, and including both with and without explicit nutrient recycling.

## References

Aguilera, G., Riggi, L., Miller, K., Roslin, T. & Bommarco, R. (2021). Organic fertilization suppresses aphid growth via carabids in the absence of specialist predators. Journal of Applied Ecology, accepted.

Allen, D.C. & Wesner, J.S. (2016). Synthesis: Comparing effects of resource and consumer fluxes into recipient food webs using meta-analysis. Ecology, 97, 11.

Allgeier, J.E., Wenger, S. & Layman, C.A. (2020). Taxonomic identity best explains variation in body nutrient stoichiometry in a diverse marine animal community. Scientific Reports, 10, 1–10.

Attayde, J.L. & Ripa, J. (2008). The Coupling Between Grazing and Detritus Food Chains and the Strength of Trophic Cascades Across a Gradient of Nutrient Enrichment. Ecosystems, 11, 980–990.

Bar-On, Y.M., Phillips, R. & Milo, R. (2018). The biomass distribution on Earth. Proceedings of the National Academy of Sciences, 115, 6506–6511.

Barabás, G., Michalska-Smith, M.J. & Allesina, S. (2017). Self-regulation and the stability of large ecological networks. Nature Ecology & Evolution, 1, 1870–1875.

Barbier, M. & Loreau, M. (2019). Pyramids and cascades: A synthesis of food chain functioning and stability. Ecology Letters, 22, 405–419.

Baxter, C.V., Fausch, K.D., Murakami, M. & Chapman, P.L. (2004). Fish invasion restructures stream and forest food webs by interrupting reciprocal prey subsidies. Ecology, 85, 2656–2663.

Berman, T. & Bronk, D. (2003). Dissolved organic nitrogen: A dynamic participant in aquatic ecosystems. Aquatic Microbial Ecology, 31, 279–305.

Bobbink, R., Hicks, K., Galloway, J., Spranger, T., Alkemade, R., Ashmore, M., Bustamante, M., Cinderby, S., Davidson, E., Dentener, F., Emmett, B., Erisman, J.W., Fenn, M., Gilliam, F., Nordin, A., Pardo, L. & De Vries, W. (2010). Global assessment of nitrogen deposition effects on terrestrial plant diversity: A synthesis. Ecological Applications, 20, 30–59.

Brockie, R.E. & Moeed, A. (1986). Animal biomass in a New Zealand forest compared with other parts of the world. Oecologia, 70, 24–34.

Buendía, C., Kleidon, A., Manzoni, S., Reu, B. & Porporato, A. (2018). Evaluating the effect of nutrient redistribution by animals on the phosphorus cycle of lowland Amazonia. Biogeosciences, 15, 279–295.

Čapek, P., Manzoni, S., Kaštovská, E., Wild, B., Diáková, K., Bárta, J., Schnecker, J., Biasi, C., Martikainen, P.J., Alves, R.J.E. et al. (2018). A plant–microbe interaction framework explaining nutrient effects on primary production. Nature ecology & evolution, 2, 1588–1596.

Cebrian, J. (1999). Patterns in the Fate of Production in Plant Communities. The American Naturalist, p. 20.

Cebrian, J. (2004). Role of first-order consumers in ecosystem carbon flow: Carbon flow through first-order consumers. Ecology Letters, 7, 232–240.

Cebrian, J. & Lartigue, J. (2004). Patterns of herbivory and decomposition in aquatic and terrestrial ecosystems. Ecological Monographs, 74, 237–259.

Cherif, M. & Loreau, M. (2009). When microbes and consumers determine the limiting nutrient of autotrophs: A theoretical analysis. Proceedings of the Royal Society B: Biological Sciences, 276, 487–497.

Darimont, C.T., Paquet, P.C. & Reimchen, T.E. (2009). Landscape heterogeneity and marine subsidy generate extensive intrapopulation niche diversity in a large terrestrial vertebrate. Journal of Animal Ecology, 78, 126–133.

De Mazancourt, C., Loreau, M. & Abbadie, L. (1998). Grazing optimization and nutrient cycling: when do herbivores enhance plant production? Ecology, 79, 2242–2252.

Elser, J.J., Fagan, W.F., Denno, R.F., Dobberfuhl, D.R., Folarin, A., Huberty, A., Interlandi, S., Kilham, S.S., McCauley, E., Schulz, K.L., Siemann, E.H. & Sterner, R.W. (2000). Nutritional constraints in terrestrial and freshwater food webs. Nature, 408, 578–580.

Galiana, N., Arnoldi, J.F., Barbier, M., Acloque, A., Mazancourt, C. & Loreau, M. (2021). Can biomass distribution across trophic levels predict trophic cascades? Ecology Letters, 24, 464–476.

Gounand, I., Mouquet, N., Canard, E., Guichard, F., Hauzy, C. & Gravel, D. (2014). The Paradox of Enrichment in Metaecosystems. The American Naturalist, 184, 752–763.

Groffman, P.M. & Rosi-Marshall, E.J. (2013). The Nitrogen Cycle. In: Fundamentals of Ecosystem Science. Elsevier, pp. 137–158.

Hagen, E.M., McCluney, K.E., Wyant, K.A., Soykan, C.U., Keller, A.C., Luttermoser, K.C., Holmes, E.J., Moore, J.C. & Sabo, J.L. (2012). A meta-analysis of the effects of detritus on primary producers and consumers in marine, freshwater, and terrestrial ecosystems. Oikos, 121, 1507–1515.

Hairston, N.G. & Hairston, N.G. (1993). Cause-Effect Relationships in Energy Flow, Trophic Structure, and Interspecific Interactions. The American Naturalist, 142, 379–411.

Hairston, N.G., Smith, F.E. & Slobodkin, L.B. (1960). Community Structure, Population Control, and Competition. The American Naturalist, 94, 421–425.

Hatton, I.A., McCann, K.S., Fryxell, J.M., Davies, T.J., Smerlak, M., Sinclair, A.R.E. & Loreau, M. (2015). The predator-prey power law: Biomass scaling across terrestrial and aquatic biomes. Science, 349, aac6284–aac6284.

Hines, J., Megonigal, J.P. & Denno, R.F. (2006). Nutrient subsidies to belowground microbes impact aboveground food web interactions. Ecology, 87, 1542–1555.

Hohener, P. & Gachter, R. (1993). Prediction of dissolved inorganic nitrogen (DIN) concentrations in deep, seasonally stratified lakes based on rates of DIN input and N removal processes. Aquatic Sciences, 55, 112–131.

Jonsson, T. (2017). Conditions for Eltonian Pyramids in Lotka-Volterra Food Chains. Scientific Reports, 7.

Leroux, S.J. & Loreau, M. (2008). Subsidy hypothesis and strength of trophic cascades across ecosystems: Subsidies and trophic cascades. Ecology Letters, 11, 1147–1156.

Loreau, M. & Holt, R.D. (2004). Spatial Flows and the Regulation of Ecosystems. The American Naturalist, 163, 606–615.

Manzoni, S., Capek, P., Porada, P., Thurner, M., Winterdahl, M., Beer, C., Briichert, V., Frouz, J., Herrmann, A.M., Lindahl, B.D. et al. (2018). Reviews and syntheses: Carbon use efficiency from organisms to ecosystems–definitions, theories, and empirical evidence. Biogeosciences, 15, 5929–5949.

Marczak, L.B., Thompson, R.M. & Richardson, J.S. (2007). Meta-analysis: Trophic level, habitat, and productivity shape the food web effects of resource subsidies. Ecology, 88, 140–148.

McCary, M.A., Phillips, J.S., Ramiadantsoa, T., Nell, L.A., McCormick, A.R. & Botsch, J.C. (2020). Transient top-down and bottom-up effects of resources pulsed to multiple trophic levels. Ecology.

Middelboe, A.L. & Markager, S. (1997). Depth limits and minimum light requirements of freshwater macrophytes. Freshwater Biology, 37, 553–568.

Montagano, L., Leroux, S.J., Giroux, M.A. & Lecomte, N. (2018). The strength of ecological subsidies across ecosystems: A latitudinal gradient of direct and indirect impacts on food webs. Ecology Letters.

Moore, J.C., Berlow, E.L., Coleman, D.C., Ruiter, P.C., Dong, Q., Hastings, A., Johnson, N.C., McCann, K.S., Melville, K., Morin, P.J., Nadelhoffer, K., Rosemond, A.D., Post, D.M., Sabo, J.L., Scow, K.M., Vanni, M.J. & Wall, D.H. (2004). Detritus, trophic dynamics and biodiversity: Detritus, trophic dynamics and biodiversity. Ecology Letters, 7, 584–600.

Nakano, S. & Murakami, M. (2001). Reciprocal subsidies: Dynamic interdependence between terrestrial and aquatic food webs. Proceedings of the National Academy of Sciences, 98, 166–170.

Newsome, T.M., Dellinger, J.A., Pavey, C.R., Ripple, W.J., Shores, C.R., Wirsing, A.J. & Dickman, C.R. (2015). The ecological effects of providing resource subsidies to predators: Resource subsidies and predators. Global Ecology and Biogeography, 24, 1–11.

Palumbi, S.R. (2003). Ecological subsidies alter the structure of marine communities. Proceedings of the National Academy of Sciences, 100, 11927–11928.

Paul, E.A. (2016). The nature and dynamics of soil organic matter: Plant inputs, microbial transformations, and organic matter stabilization. Soil Biology and Biochemistry, 98, 109–126.

Perkins, M.J., Inger, R., Bearhop, S. & Sanders, D. (2018). Multichannel feeding by spider functional groups is driven by feeding strategies and resource availability. Oikos, 127, 23–33.

Polis, G.A., Anderson, W.B. & Holt, R.D. (1997). Toward an integration of landscape and food web ecology: The dynamics of spatially subsidized food webs. Annual Review of Ecology and Systematics, 28, 289–316.

Polis, G.A., Myers, C.A. & Holt, R.D. (1989). The Ecology and Evolution of Intraguild Predation: Potential Competitors That Eat Each Other. Annual Review of Ecology and Systematics, 20, 297–330.

Raun, W.R. & Johnson, G.V. (1999). Improving Nitrogen Use Efficiency for Cereal Production. Agronomy Journal, 91, 357–363.

Riggi, L.G. & Bommarco, R. (2019). Subsidy type and quality determine direction and strength of trophic cascades in arthropod food web in agro-ecosystems. Journal of Applied Ecology.

Rooney, N., McCann, K., Gellner, G. & Moore, J.C. (2006). Structural asymmetry and the stability of diverse food webs. Nature, 442, 265–269.

Scaini, A., Zamora, D., Livsey, J., Lyon, S.W., Bommarco, R., Weih, M., Jaramillo, F. & Manzoni, S. (2020). Hydro-climatic controls explain variations in catchmentscale nitrogen use efficiency. Environmental Research Letters, 15, 094006.

Shurin, J.B., Gruner, D.S. & Hillebrand, H. (2006). All wet or dried up? Real differences between aquatic and terrestrial food webs. Proceedings of the Royal Society B: Biological Sciences, 273, 1–9.

Spiecker, B., Gouhier, T.C. & Guichard, F. (2016). Reciprocal feedbacks between spatial subsidies and reserve networks in coral reef meta-ecosystems. Ecological Applications, 26, 264–278.

Wollrab, S., Diehl, S. & De Roos, A.M. (2012). Simple rules describe bottom-up and top-down control in food webs with alternative energy pathways. Ecology Letters, 15, 935–946.

Zou, K., Thébault, E., Lacroix, G. & Barot, S. (2016). Interactions between the green and brown food web determine ecosystem functioning. Functional Ecology, 30, 1454–1465.

